# A versatile Halo- and SNAP-tagged BMP/TGFβ receptor library for quantification of cell surface ligand binding

**DOI:** 10.1101/2022.03.04.482944

**Authors:** Jerome Jatzlau, Wiktor Burdzinski, Michael Trumpp, Leon Obendorf, Kilian Roßmann, Katharina Ravn, Marko Hyvönen, Francesca Bottanelli, Johannes Broichhagen, Petra Knaus

**Affiliations:** Institute of Chemistry and Biochemistry - Biochemistry, Berlin, Germany; Berlin-Brandenburg School for Regenerative Therapies (BSRT), Berlin, Germany; Leibniz-Forschungsinstitut für Molekulare Pharmakologie, Berlin, Germany; Department of Biochemistry, University of Cambridge, Cambridge, UK

**Author notes:** Equal contributing authors (Jerome Jatzlau, Wiktor Burdzinski).

## Abstract

The TGFβ superfamily of secreted growth factors comprises more than 30 members including TGFβs, BMPs and Activins. While all TGFβ superfamily members signal through heteromeric receptor complexes to regulate a plethora of developmental and homeostatic processes, each ligand possesses a unique affinity towards a subset of BMP and TGFβ type I and type II receptors. Whereas the Activin and TGFβ class display a higher affinity towards type II receptors, BMPs and GDFs preferentially bind to type I receptors. Sofar, the lack of specific antibodies and chemical biology tools hampered simultaneous testing of ligand binding towards all BMP and TGFβ receptors. Here we present a N-terminally Halo- and SNAP-tagged TGFβ/BMP receptor library to visualize the receptor complexes in dual color. In combination with novel fluorescently labeled TGFβ superfamily ligands, we established a Ligand Surface Binding Assay (LSBA) for optical quantification of receptor-dependent growth factor binding for Activin A, TGFβ1 and BMP9 in a cellular context. We confirm ligand-receptor interface specificity by identifying BMPR2- or ALK2-mutants that switch from a low-affinity Activin A- or BMP9-receptor to a high-affinity receptor, respectively.

## Introduction

The transforming growth factor β (TGFβ) family compromises more than 30 ligands, including bone morphogenetic proteins (BMPs) and activins, which regulate a broad range of developmental and homeostatic functions in a variety of cell types and organs, including the osteochondral and the vascular system (Hiepen, Jatzlau, & Knaus, 2020; Hiepen, Yadin, Rikeit, Dorpholz, & Knaus, 2016; Nickel & Mueller, 2019; Nickel, Ten Dijke, & Mueller, 2018; Yadin, Knaus, & Mueller, 2016). The clinical impact of the TGFβ/BMP signalling pathways is highlighted by numerous examples, in which dysbalanced SMAD signalling as a result of genetic mutations is causative for several vascular diseases such as pulmonary arterial hypertension (PAH), hereditary hemorrhagic telangiectasia (HHT) or aortic aneurysm but also for the devastating musculoskeletal disease fibrodysplasia ossificans progressiva (FOP) (Morse, Deng, & Knowles, 2001; Shore et al., 2007; Shovlin, 1997). Taking into account their diverse involvement in physiological and pathological processes, BMPs are increasingly referred to as body rather than bone morphogenetic proteins (Wagner et al., 2010). TGFβ and BMP signal transduction is initiated by oligomerization of two type I and two type II serine/threonine kinase receptors upon binding one dimeric ligand. Trans-phosphorylation of the GS-box of a type I receptor by a type II receptor renders the type I receptor active, thereby allowing the recruitment and activation of SMAD proteins which act as transcriptional regulators (Nickel & Mueller, 2019; Nickel et al., 2018).

TGFβ superfamily ligands exhibit preferential binding to either type I or type II receptors. BMPs and some growth and differentiation factors (GDFs) were described to possess high affinity towards type I receptors, which induce SMAD1/5/8 signalling, e.g. BMP9 for ALK1, BMP6 for ALK2, BMP2 for ALK3 and GDF5 for ALK6 (David, Mallet, Mazerbourg, Feige, & Bailly, 2007; Heinecke et al., 2009; Nishitoh et al., 1996; Salmon et al., 2020). In contrast SMAD2/3 activating TGFβ superfamily members exhibit high affinity towards type II receptors, e.g. TGFβ1 for TGFBR2 or Activin A, GDF8 and GDF11 for ACVR2B (Goebel et al., 2022; Heinecke et al., 2009; Rodriguez, Chen, Weinberg, & Lodish, 1995; Walker et al., 2017). However, some receptors are promiscuous in ligand binding (e.g. ACVR2B) and receptor expression levels vary in specific cell types. When high-affinity receptors are absent this often results in cell type specific ligand competition scenarios between low-affinity receptors (Martinez-Hackert, Sundan, & Holien, 2021). This is particularly important in pathophysiological conditions with unbalanced BMP signalling, as recently shown for example for PAH, in which the absence of BMPR2 increases responsiveness towards TGFβ1 (Hiepen et al., 2019).

All TGFβ family members are dimers that resemble two left hands with palms together, forming a butterfly-like structure. Together both hands offer two ‘wrist’ and two ‘knuckle’ epitopes, which bind to type I and type II receptors, respectively (Yadin et al., 2016). Indeed, structural variation in these ligands yields different binding modes of type I and type II receptors for the BMP, TGFβ and Activin classes (Goebel, Corpina, et al., 2019; Martinez-Hackert et al., 2021). While both receptor types bind independently BMPs and Activins, TGFβ receptors bind cooperatively to TGFβs. Hereby TGFBR2 binds uniquely to the fingertips of TGFβ1-3, thus creating a shared epitope with the ligands for the low-affinity type I receptor ALK5 (i.e. cooperative binding) (Groppe et al., 2008). In contrast, ACVR2A, ACVR2B and likely also BMPR2 bind via hydrophobic interactions to the knuckle epitope (Kirsch, Nickel, & Sebald, 2000; Thompson, Woodruff, & Jardetzky, 2003; Townson et al., 2012). High-affinity binding of BMP type I receptors to the wrist epitope can be attributed to a structurally unique α1 helix, which projects a phenylalanine into a hydrophobic pocket of the ligands (ALK3 and ALK6) or forms polar bonds with both ligand monomers (ALK1), in a so-called lock and key mechanism (Goebel, Hart, McCoy, & Thompson, 2019; Kirsch, Sebald, & Dreyer, 2000; Kotzsch, Nickel, Sebald, & Mueller, 2009; Townson et al., 2012). Finally, Activins possess significant variability in the dimeric structure and are thought to become restrained upon type II receptor binding, adopting a conformation that together with the fingertips selects for type I receptor recruitment (i.e. conformational selection) (Goebel, Corpina, et al., 2019; Goebel et al., 2022; Greenwald et al., 2004; Thompson et al., 2003).

Whereas co-crystallization of the ligands with their respective receptor extracellular domains (ECD) but also extensive surface plasmon resonance (SPR) experiments provided the structural framework and *in vitro* ligand receptor affinities, a spatiotemporal resolution of the complexes with full-length membrane bound receptors could so far only be achieved via microscopic approaches. Using immunofluorescence co-patching of epitope-tagged BMP receptors allowed the identification of preformed receptor complexes (PFCs) and BMP-induced signaling complexes (BISCs) (Nohe et al., 2002). The dynamic association of BMP receptors into homo- and heterodimers and the resulting lateral diffusion was shown by fluorescence recovery after photobleaching (FRAP) and Patch/FRAP experiments (Marom, Heining, Knaus, & Henis, 2011; Nohe et al., 2002). The dynamics of the short-lived BMP-induced heteromeric receptor complex formation were investigated by using two-color single particle tracking (SPT) utilizing quantum dots (QDs) (Guzman et al., 2012). However, most of the previous approaches relied on antibody recognition of epitope-tagged receptors, resulting in unknown fluorophore stoichiometries, requiring long incubation times and restricting analysis to fixed cells. To overcome this, tagging receptors with self-labeling enzymes would allow for live time analysis using straightforward staining protocols. Self-labeling protein tags like the SNAP-(derived from mammalian O^6^-alkylguanine DNA alkyl transferase) or the Halo-tag (derived from bacterial haloalkane dehalogenase) specifically react with benzyl guanine (BG) or chloroalkane (CA), respectively, transferring one fluorophore of choice by means of covalent linkage, enabling fast and stoichiometric antibody-free labeling of a protein of interest (POI) (Juillerat et al., 2003; Los et al., 2008). Simultaneous application of both self-labeling protein tags for analysis of two POIs previously demonstrated a two-color visualization approach for intracellular targets (Bottanelli et al., 2016). Addition of sulfonate groups to SNAP- and Halo-tag ligands renders them cell-impermeable and thereby allows discrimination of receptor populations by exclusively staining cell surface exposed receptors (Birke et al., 2022; Poc et al., 2020). A step into this direction has been reported in a study where two N-terminally SNAP-tagged receptors have been used for FRAP analysis, namely BMPR2 and ALK3 (Mundy, Yang, Takano, Billings, & Pacifici, 2018).

In order to visualize all TGFβ/BMP receptors including their complexes in dual-color we generated a comprehensive N-terminally Halo- and SNAP-tagged receptor library. Additionally, designing and applying fluorescently labeled growth factor ligands allowed us to not only monitor receptor-dependent ligand surface binding but also to establish a Ligand Surface Binding Assay (LSBA) for screening of ligand binding under native conditions. As LSBA was suitable to test TGFβ superfamily ligand-receptor binding in living cells, this allowed us to provide additional proof for ligand-receptor interface specificities. To provide additional proof for our ligand-receptor interaction analyses, we designed and generated variants of type I and type II receptors with altered ligand specificities, as verified by microscopy. This work provides a new adaptable visualization platform for color multiplexing, suitable for super resolution microscopy and allowing to investigate TGFβ/BMP ligand specificities to their respective receptor in living cells.

## Results

### 1. Cell surface staining of Halo- and SNAP-tagged BMP-, Activin- and TGFβ-receptors

BMPs, TGFβs and Activins signal through binding towards different combinations of BMP/TGFβ type I and type II receptors in a heterotetrameric receptor ligand complex (**Fig. 1A**). Hereby the type I and the type II receptor ECD bind to the wrist and knuckle epitope of TGFβ family ligands, respectively (**Fig. 1B**). We here aim to visualize preferentially ligand receptor binding on living cells using state of the art imaging tools. Limited visualization of BMP and TGFβ receptors by immunofluorescence can be overcome by utilization of self-labeling Halo- and/or SNAP-tag enzymes. BMP/TGFβ receptors, including type I receptors (hALK1-ALK6, rALK7 and hALK2-R206H) and type II receptors (hACVR2A, hACVR2B, hBMPR2 and hTGFBR2) were therefore N-terminally fused with Halo- or SNAP-tags separated by a 5x glycine linker downstream the signal peptide sequence, to obtain the receptor library shown in (**Fig. 1C**). All Halo- and SNAP-tagged type I and type II receptor constructs were transiently expressed in COS-7 cells and lysates were subjected to Western blot analysis with specific antibodies directed against Halo- (**Fig. S1A**) or SNAP-tags (**Fig. S1B**). While untagged type I receptors have molecular weights between approx. 56-60 kDa, the Halo- or SNAP-tagged type I receptors were detected at a molecular weight corresponding to the addition of a Halo-tag (^~^33 kDa) or a SNAP-tag (^~^20 kDa) (**Fig. S1A,B**). Similar weight shifts were observed for Halo- and SNAP-tagged type II receptors BMPR2, ACVR2A, ACVR2B and TGFBR2 (**Fig S1A,B**). Further, all Halo- and SNAP-tagged receptors were positively detected using impermeable CA-Alexa488 or BG-Alexa488 staining, respectively. Using cell impermeable dyes provides the advantage to visualize exclusively the surface receptor pool, while cell permeable dyes or fluorescent protein tags (e.g. GFPs) would stain both the membrane and intracellular fraction. As an example, we stained COS-7 cells transiently expressing ALK1-SNAP with first cell-impermeable SNAP-tag substrate BG-ATTO594, chased by cell-permeable BG-SiR, highlighting the advantage of surface-only staining (**Fig. 1D**). ATTO594 uniquely stained receptors located at the plasma membrane while SiR visualizes the crowded intracellular fraction. With the additional advantage that STED compatible dyes were used, super-resolution imaging allowed a lateral resolution of ^~^ 40 nm pixel (**Fig. S1C**). Together, this describes a full library of BMP and TGFβ receptors and showcases that CA- and BG-coupled dyes allow to exclusively study the cell surface population by STED nanoscopy.

**Figure 1.**
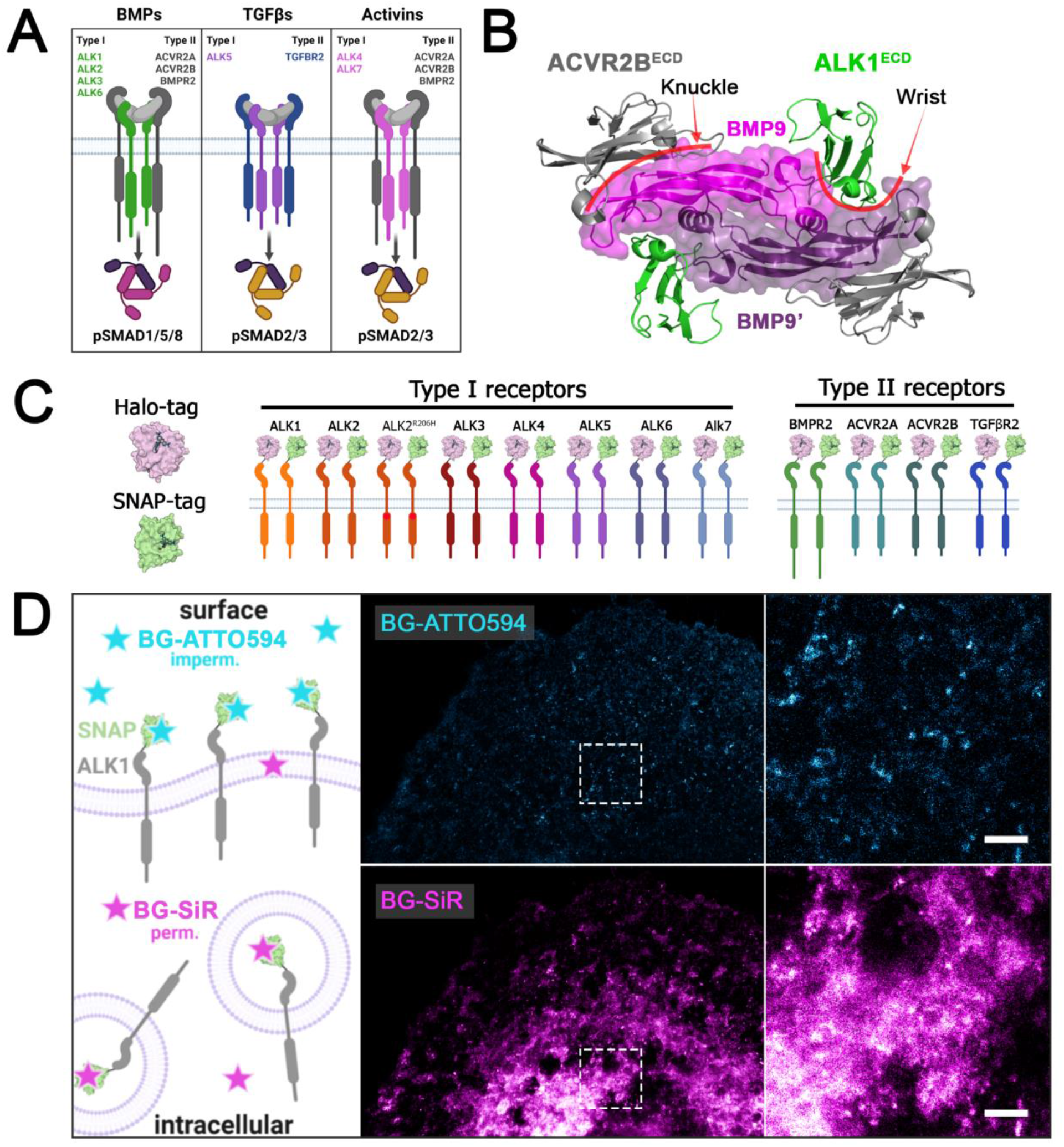
Halo-& SNAP-tagged BMP and TGFβ receptors can be visualized covalently and fast with organic fluorophore-labelled substrates allowing the discrimination between surface and intracellular receptor populations: **(A)** Overview of preferential receptor-ligand paring. **(B)** Top view of the extracellular domains of ALK1 and ACVR2B in a heterotetrameric receptor complex bound to BMP9 (crystal structure PDB: 4FAO). Type l receptor ALK1 binds at the wrist interface, type II receptor ACVR2B binds at the knuckle interface of BMP9, respectively. **(C)** Schematic of N-terminally Halo- and SNAP-tagged BMP and TGFβ receptor library consisting of type I receptors (ALK1-ALK7 and FOP-mutant ALK2-R206H) and type II receptors (BMPR2, ACVR2A, ACVR2B and TGFBR2). **(D)** Discrimination between surface and cytosolic receptor populations. COS-7 cells transiently expressing ALK1-SNAP were 24 hours post transfection incubated with BG-ATTO594 (impermeable; cyan) and chased by BG-SiR (permeable; magenta) allowing for staining of the surface receptor population and the cytosolic receptor population, respectively. (BG: benzyl guanine; SiR: silicon rhodamine).

### 2. High-affinity binding of Activin A by BMP type II receptors

Specificity of Activin A binding to BMP/TGFβ receptors was previously mainly studied in cell-free SPR measurements utilizing immobilized receptor ECDs and soluble Activin A (Greenwald et al., 2004; Thompson et al., 2003). Binding and chemical crosslinking of ^125^I-labeled Activin A in transfected COS-7 cells (Attisano, Wrana, Cheifetz, & Massague, 1992; Donaldson, Mathews, & Vale, 1992) allowed for the detection of ligand bound receptors after cell lysis and by radiography. Here, we aimed to visualize receptor dependent Activin A binding on living transfected COS-7 cells in a fluorescence-based approach. Fluorescently labeled Activin A was recently used to show ACVR2B clustering (Ramachandran et al., 2021). Accordingly, we have labeled Activin A with Cy5 and established an analysis pipeline for assessment of Activin A-Cy5 binding on COS-7 cells dependent on the expression of various Halo-tagged receptors (**Fig. 2A**). Receptors from the Halo-tagged BMP/TGFβ receptor library (**Fig. 1C**) were transiently expressed in COS-7 cells and stained with cell-impermeable Halo-tag substrate CA-Alexa488 at 4 °C to impede internalization. Simultaneous incubation with Activin A-Cy5 allowed for visualization of ligand binding to the respective receptor. Acquired confocal images were semi-automatically analyzed with Fiji ImageJ to quantify Activin A-Cy5 and receptor signal intensities (**Fig. 2A**). As expected ACVR2B led to the highest (^~^84-fold) Activin A binding, followed by ACVR2A (^~^29-fold) and BMPR2 (^~^27-fold), whereas TGFBR2 and all type I receptors showed no significant increase in Activin A binding (**Fig. 2B-C, Fig. S2A-B**). Further, Activin A-Cy5 and receptor intensities positively correlated for ACVR2A, ACVR2B and BMPR2, while TGFBR2 expression did not correlate with ligand binding, highlighting that increased high-affinity receptor expression leads also to more ligand binding (**Fig. 2D**). Live cell imaging (LCI) of COS-7 cells expressing ACVR2B-Halo highlighted that Activin A-Cy5 binding occurs already within minutes after ligand addition (**Movie 1** & **Fig. S3**). Overall, these results are in accordance with binding and cross-linking studies in COS cells utilizing ^125^I-Activin A (Attisano et al., 1992), as well as SPR studies using immobilized receptors (Heinecke et al., 2009) but reflect binding on living cells and under native conditions.

**Figure 2.**
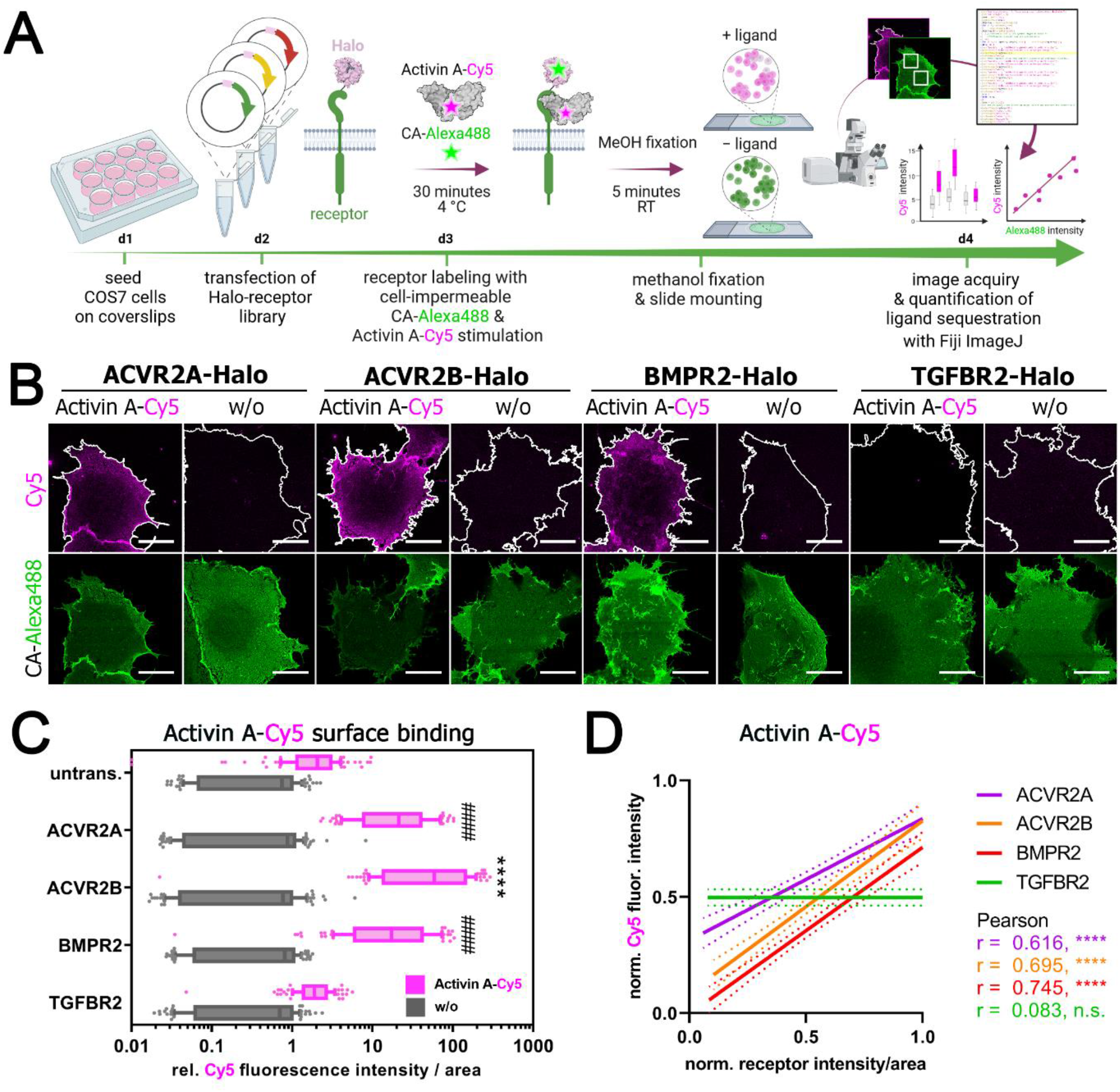
Activin A-Cy5 binding by BMP type II receptors: **(A)** Schematic illustration for visualization of fluorescent growth factor binding on COS-7 cells expressing Halo-tagged BMP or TGFβ receptor constructs. **(B-D)** COS-7 cells were seeded on coverslips and transfected with indicated Halo-tagged receptor constructs. 24 hours post transfection, cells were incubated with non-permeable fluorescent Halo-tag substrate CA-Alexa488 (green) and Activin-Cy5 (magenta) for 30 minutes at 4 °C, fixated with methanol for 5 minutes at room temperature and mounted on glass slides. Cells were imaged at a confocal microscope and 10 cells per condition and replicate were analysed with a semi-automated Fiji ImageJ macro pipeline for assessment of fluorescent growth factor binding (Activin A-Cy5) and fluorescence intensity of receptors (CA-Alexa488). Four ROIs of 100 μm^2^ were quantified in each cell. (n = 3 independent experiments) **(B)** Representative confocal microscopy images of COS-7 cells transiently expressing ACVR2A-, ACVR2B-, BMPR2- or TGFBR2-Halo incubated with CA-Alexa488 and simultaneously stimulated with Activin A-Cy5 or PBS as control. Scale bar ≙ 20 μm. **(C)** Activin A-Cy5 surface binding is represented as relative fluorescence intensity per area. Data is shown as F.I. ± SD. Significance was calculated using two-way ANOVA and Tukey’s post-hoc test. #### p < 0.0001 ≡ significant relative to Activin A-Cy5 stimulated control cells, ****p < 0.0001 ≡ significant relative to all Activin A-Cy5 stimulated conditions including ACVR2A and BMPR2. **(D)** Linear regression and correlation analysis of ligand:receptor binding based on Cy5-fluorescence intensity and normalized receptor fluorescence (CA-Alexa488) per area. (CA: chloroalkane).

### 3. SiR-d12 labelled-BMP9 and - TGFβ1 allow for the identification of their respective high-affinity receptors

To extend the studies to other members of the TGFβ ligand super-family, we labelled BMP9 and TGFβ1 using *N*-hydroxysuccinimide (NHS) silicon rhodamine-d12 (SiR-d12) esters (Nanda & Lorsch, 2014; Roßmann et al., 2020) (**Fig. S4**) to obtain far-red fluorescent BMP9-SiR-d12 and TGFβ-SiR-d12 (**Fig. 3A; center**). The labelling protocol is described in the method section and a scheme of SiR-d12 labelled ligands binding to their respective high-affinity Halo-tagged receptors and staining parameters are depicted in (**Fig. 3A; left & right**). We next tested TGFβ1-SiR-d12 and BMP9-SiR-d12 binding on COS-7 cells, transiently expressing receptors found in signalling complexes of the respective ligand, i.e. TGFβ1-TGFBR2-ALK5 and BMP9-ACVR2B-ALK1 (Radaev et al., 2010; Townson et al., 2012). As expected, we observed strong TGFβ1-SiR-d12 binding to cells expressing TGFBR2 but not to cells expressing ALK5 (**Fig. 3B**). Similarly, the BMP9-high affinity receptor ALK1 strongly increased binding of BMP9-SiR-d12, while ACVR2B, which was shown by SPR to exhibit a similar high picomolar affinity (ALK1-Fc K_D_ = 31.3 pM, ACVR2B K_D_ = 33.0 pM)(Townson et al., 2012), failed in binding BMP9-SiR-d12 (**Fig. 3B-C**). Both TGFβ1-SiR-d12 and BMP9-SiR-d12 signal intensity positively correlated to TGFBR2 or ALK1 receptor intensity, respectively (**Fig. 3D**). Finally, we expanded our ligand surface binding assay to the whole receptor library but did not observe any other significant increases for neither TGFβ1-SiR-d12 nor BMP9-SiR-d12 binding by any other receptor (**Fig. 3C-D, Fig.S5 & S6**).

**Figure 3.**
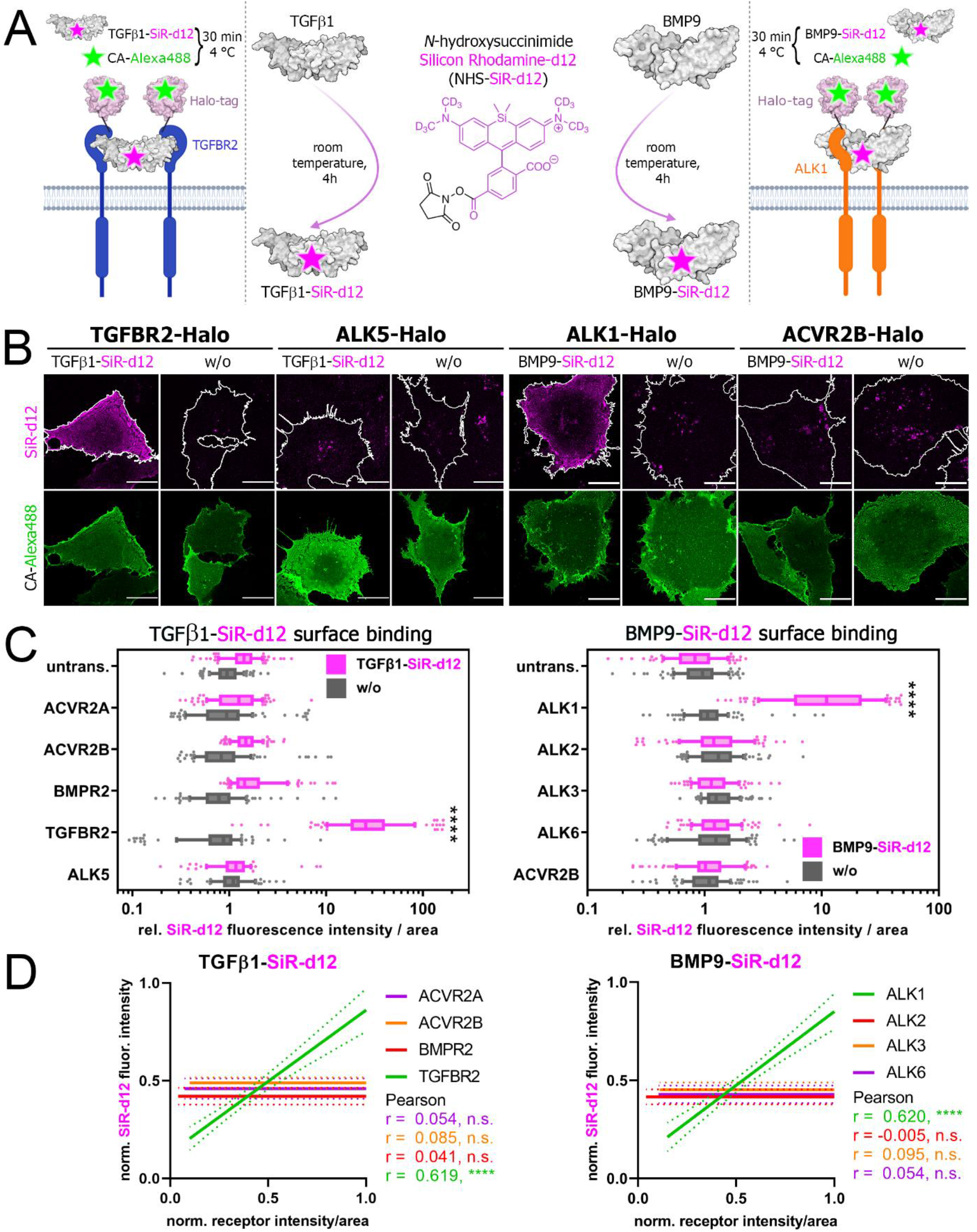
Silicon Rhodamine-d12 labelling of TGFβ1 and BMP9 visualizes binding by TGFBR2 and ALK1, respectively: **(A-D)** Transiently transfected COS-7 cells expressing TGFBR2-, ALK5-, ALK1- or ACVR2B-Halo were 24 hours post transfection simultaneously incubated with Halo-tag substrate CA-Alexa488 (green) and TGFβ1-SiR-d12 or BMP9-SiR-d12 (magenta). **(A)** Ligand labelling strategy of TGFβ1 and BMP9 utilizing *N*-hydroxysuccinimide deuterated silicon rhodamine ester (NHS-SiR-d12) (middle). Illustration of TGFβ1-SiR-d12 and BMP9-SiR-d12 binding to their respective high-affinity receptor TGFBR2 (left) or ALK1 (right). **(B)** Representative confocal microscopy images of TGFβ1-SiR-d12 (left) and BMP9-SiR-d12 stimulated COS-7 cells expressing respective high-(TGFBR2, ALK1) or low-affinity receptors (ALK5, ACVR2B). Scale bar ≙ 20 μm. **(C)** TGFβ1-SiR-d12 (left) and BMP9-SiR-d12 (right) surface binding represented as relative fluorescence intensity per area. Data is shown as F.I. ± SD. Significance was calculated using two-way ANOVA and Tukey’s post-hoc test. ****p < 0.0001 ≡ significant relative to all stimulated conditions, (n = 3 independent experiments). **(D)** Linear regression and correlation analysis of ligand:receptor binding based on SiR-d12-fluorescence intensity and normalized receptor fluorescence (CA-Alexa488) per area (n = 3). (CA: chloroalkane).

### 4. Receptor-mimics reveal critical residues for Activin A-type II receptor interfaces

Binding of Activin A-Cy5 to cell surface expressed type II receptors confirmed a higher affinity of ACVR2B over ACVR2A and BMPR2, which showed comparable ligand binding (**Fig. 2**) (Heinecke et al., 2009). Structural analysis of Activin A bound to ACVR2B previously highlighted that the core of the concave ACVR2B interface contains hydrophobic residues, which interact with hydrophobic residues in the Activin A knuckle epitope, surrounded by polar residues at the edge that form strong electrostatic interactions (Thompson et al., 2003) (**Fig. 4A**). Sequence comparison of ACVR2B with ACVR2A indicated only subtle differences in two amino acids which form hydrophobic bonds with Activin A (Tyr_A2B_60/Phe_A2A_61, Phe_A2B_82/IleA2A83), whereas BMPR2 is characterized by a different elongated 2/3 loop, which harbors mostly polar and charged amino acids of the Activin A epitope in ACVR2B (Leu_A2B_79-Asp_A2B_80-Asp_A2B_81-Phe_A2B_82-Asn_A2B_83) (**Fig. 4A**). Evolutionary conserved residues and potentially crucial for ligand binding properties in this area include Trp78 (**Fig. 4A**). We next asked if substituting half of the BMPR2 2/3 loop (Ser _BR2_84-X-Gln_BR2_90) with the Activin A-binding ACVR2B amino acids (Leu_A2B_79-X-Asn_A2B_83) would increase ligand binding of BMPR2. The resulting BMPR2^Mimic-ACVR2B^-Halo construct was generated (**Fig. S7A**) and Activin A-Cy5 binding was analyzed as described above (**Fig. 2A**). Indeed, BMPR2^Mimic-ACVR2B^ showed increased binding compared to BMPR2 and even to ACVR2B, highlighting the crucial role of the 2/3 loop in Activin A ligand binding (**Fig. 4C-D**). We next asked, whether the altered 2/3 loop in BMPR2^Mimic-ACVR2B^ forms similar interactions with Activin A residues as ACVR2B. We therefore, took advantage of the ACVR2B/Activin A crystal structure (PDB: 1S4Y) (Greenwald et al., 2004) and the crystallized ECD of BMPR2 (PDB: 2HLQ) (Mace, Cutfield, & Cutfield, 2006) and performed *in silico* docking of the BMPR2 structure to Activin A (**Fig. S8**). Next, we generated a homology model of BMPR2^Mimic-ACVR2B^ docked on to Activin A. Indeed, our predictions highlight that BMPR2^Mimic-ACVR2B^ forms the same electrostatic interactions as ACVR2B with Activin A at its 2/3 loop (e.g. a salt bridge between Asp_A2B_80 with Arg_ActA_87) (**Fig. 4E**), while in BMPR2 these interactions are altered (e.g. Pro_BR2_91 forms a less favorable interaction with Arg_ActA_87). The increased Activin A binding of BMPR2^Mimic-ACVR2B^ when compared to ACVR2B can be explained by the extended finger 3, present in BMPR2 and likewise in BMPR2^Mimic-ACVR2B^, which allows for additional electrostatic interactions with fingertips 1 and 2 of Activin A (e.g., Glu_BR2_109 with Asp_ActA_26) (Fig. S6C). Collectively this data highlighted that our Halo-tagged receptor library is suitable for assessment of ECD mutation screening and that the substitution of the 2/3 loop of BMPR2 with the ACVR2B 2/3 loop yields an Activin A high-affinity BMPR2.

**Figure 4.**
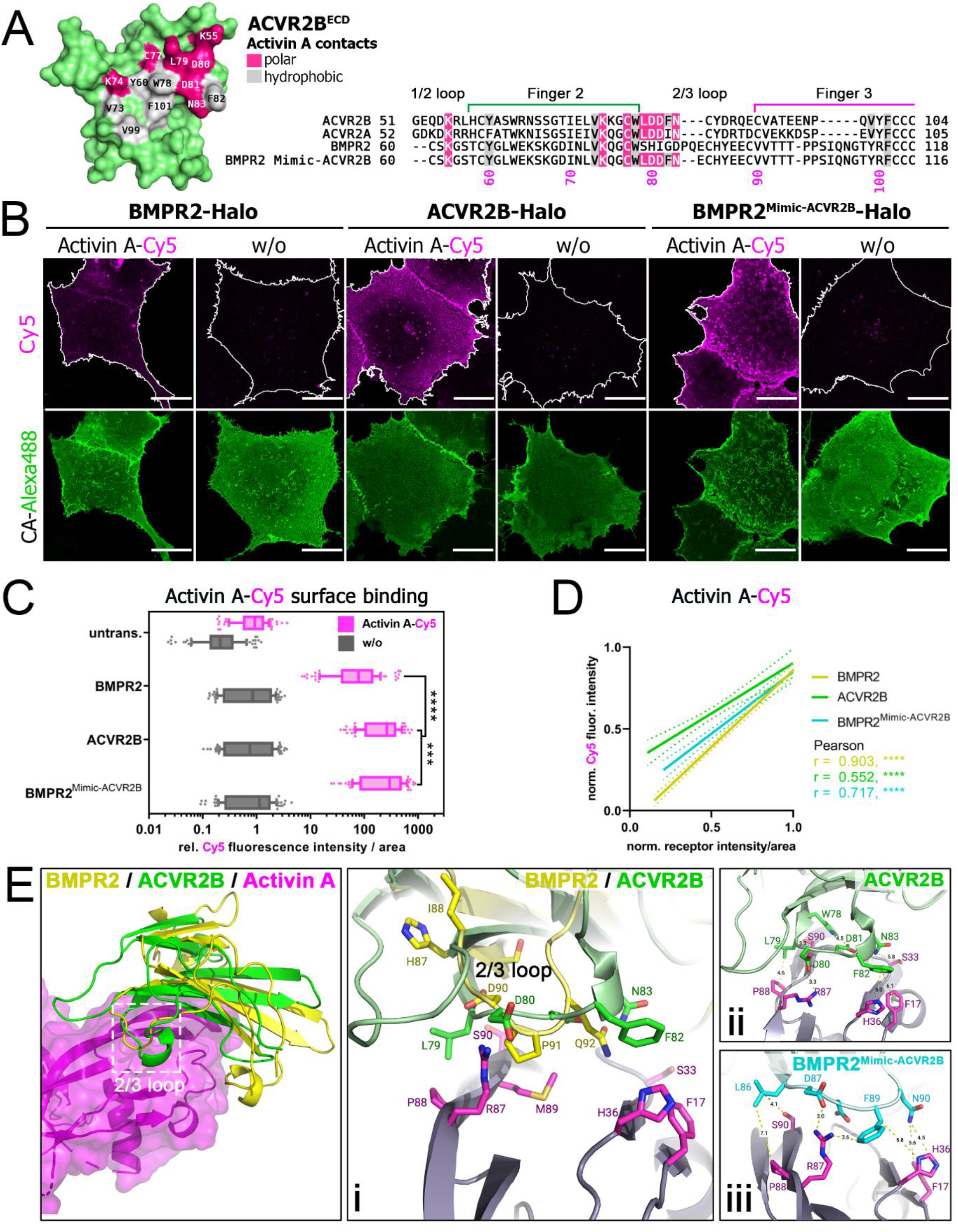
BMPR2 gains Activin A affinity through substitution of ACVR2B 2/3 loop. **(A)** Molecular surface of ACVR2B^ECD^ showing the Activin A interface (PDB: 1S4Y). Residues forming polar or hydrophobic contacts are coloured in pink and grey, respectively (left). Sequence alignment of BMP type II receptors with Activin A contact residues highlighted. BMPR2^Mimic-ACVR2B^ sequence includes ACVR2B substitutions in the 2/3 loop. **(B-D)** Transiently transfected COS-7 cells expressing BMPR2-, ACVR2B- or BMPR2^Mimic-ACVR2B^-Halo receptors were 24 hours post transfection simultaneously incubated with Halo-tag substrate CA-Alexa488 (green) and Activin A-Cy5 (magenta). (n = 3 independent experiments). **(B)** Representative confocal microscopy images of COS-7 cells transiently expressing BMPR2-, ACVR2B-or BMPR2^Mimic-ACVR2B^-Halo incubated with CA-Alexa488 and simultaneously stimulated with Activin A-Cy5 or PBS as control. Scale bar ≙ 20 μm. **(C)** Activin A-Cy5 surface binding represented as relative fluorescence intensity per area. Data is shown as F.I. ± SD. Significance was calculated using two-way ANOVA and Tukey’s post-hoc test. ***p < 0.001 ***, p < 0.0001 ≡significance as indicated (n = 3). **(D)** Linear regression and correlation analysis of ligand:receptor binding based on Cy5-fluorescence intensity and normalized receptor fluorescence (CA-Alexa488) per area (n = 3). Overview (left) and cartoon/stick representation (i) of the binding interface of the ACVR2B/Activin A (PDB: 1S4Y) crystal structure compared to our *in silico* predicted BMPR2/Activin A complex (structures derived from PDB: 2HLQ and 1S4Y, respectively). Distance measurements (using PyMOL) in Angstrom of the 2/3 loop residues of ACVR2B (ii) and BMPR2^Mimic-ACVR2B^ variant (iii) to Activin A reveal possible interactions. Docking predictions were carried out using the Rosetta docking protocols.

### 5. Generation of a BMP9-binding ALK2 receptor

After successfully increasing the affinity of a type II receptor (**Fig. 4**), we aimed to equally enhance binding of BMP9-SiR-d12 to type I receptors. For this, we selected ALK2-Halo to substitute key residues from the high affinity receptor ALK1. In line with our data (**Fig 3B,C**), ALK1 is the high affinity receptor for BMP9 (David et al., 2007; Townson et al., 2012; van Meeteren et al., 2012) and presents with a ^~^1800 times higher affinity (ALK1-Fc K_D_ = 45.5 pM) when compared to ALK2 (ALK2-Fc K_D_ = 83 nM) (Salmon et al., 2020). However, ALK2 was shown to transmit BMP9 signals in the absence of ALK1 in non-endothelial cells (Kang et al., 2004; Luo et al., 2010; Olsen et al., 2014; Salmon et al., 2020). The ALK1-BMP9 interface is subdivided into three sites (I-III). Site I and III are overlapping with other type I receptor-BMP interfaces while site II is unique to ALK1 for BMP9 and BMP10 binding (Salmon et al., 2020; Townson et al., 2012). Alongside, the helix α1 and loop F3 of ALK1 contain charged residues (**Fig. 5A**), which through polar interactions convey binding to both BMP9 monomers within the wrist epitope (Salmon et al., 2020; Townson et al., 2012). These polar residues (HERR motif - His73, Glu75, Arg78 and Arg80) bridge to amino acids that are unique to BMP9 and BMP10, thereby defining ligand specificity (Salmon et al., 2020). Sequence comparison with ALK2 highlighted that the HERR motif surrounding the helix α1 is absent in ALK2, alongside other critical residues that define ALK1/BMP9 specificity (**Fig. 5A**). We therefore generated three ALK2 variants, in which we sequentially substituted amino acids by the ALK1/BMP9 epitope (V1-3 ALK2^Mimic-ALK1^) and tested BMP9-SiR-d12 surface binding in transfected COS-7 cells (**Fig. 5B-D, Fig. S9**). Interestingly, substitution of the HERR motif only was not sufficient to increase BMP9-binding towards ALK2 (V1 ALK2^Mimic-ALK1^), however additional substitution of hydrophobic residues surrounding the HERR motif in the helix α1 rendered ALK2 BMP9-sensitive (V2 ALK2^Mimic-ALK1^). Additional substitution of amino acids at the site I interface (VVFREE-motif, V3 ALK2^Mimic-ALK1^) did not further increase ligand binding.

**Figure 5.**
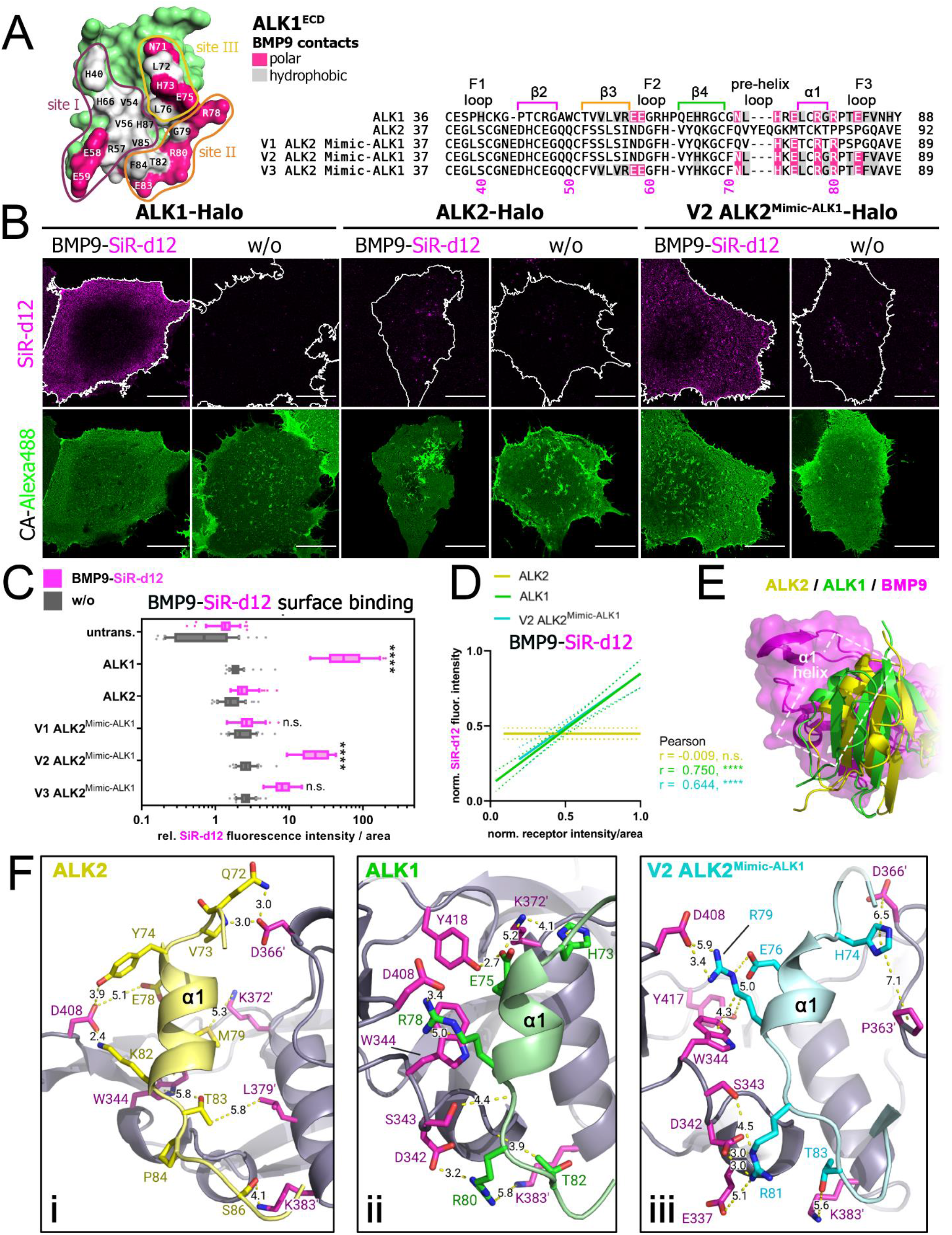
ALK2 gains BMP9 affinity through substitution of ALK1 helix α1. **(A)** Molecular surface of ALK1^ECD^ showing the BMP9 interface (PDB: 4FAO). Residues forming polar or hydrophobic contacts are coloured in pink and grey, respectively (left). Interaction sites I-III are indicated with coloured lines. Sequence alignment of ALK1, ALK2 and variants V1-3 of ALK2^Mimic-ALK1^ with BMP9 contact residues highlighted. ALK2^Mimic-ALK1^ sequences include sequential ALK1 substitutions at interaction sites II & III (V1-2) and site I (V3). **(B-D)** Transiently transfected COS-7 cells expressing ALK1-, ALK2-, V1-V3 ALK2^Mimic-ALK1^-Halo receptors were 24 hours post transfection simultaneously incubated with Halo-tag substrate CA-Alexa488 (green) and BMP9-SiR-d12 (magenta). **(B)** Representative confocal microscopy images of COS-7 cells transiently expressing ALK1-, ALK2- or V2 ALK2^Mimic-ALK1-^ Halo incubated with CA-Alexa488 and simultaneously stimulated with BMP9-SiR-d12 or PBS as control. Scale bar ≙ 20 μm. **(C)**BMP9-SiR-d12 surface binding represented as relative fluorescence intensity per area. Data is shown as F.I. ± SD. Significance was calculated using two-way ANOVA and Tukey’s post-hoc test. ****p < 0.0001 ≡ significant relative to BMP9-SiR-d12 stimulated control cells. **(D)** Linear regression and correlation analysis of ligand:receptor binding based on SiR-d12-fluorescence intensity and normalized receptor fluorescence (CA-Alexa488) per area. (**E**) Cartoon and stick representation of the “HERR”-motif residues (α1 helix and F3 Loop) of ALK2 (based on an AlphaFold model docked to BMP9 (PDB: 4FAO) using Rosetta docking protocols) (i), ALK1 co-crystallized to BMP9 (PDB: 4FAO) (ii), and V2 ALK2^Mimic-ALK1^ (iii) (derived from ALK2, mutated, and docked to BMP9 (PDB: 4FAO) using the Rosetta Commons Modelling Suite). ALK2 cannot form interactions originating from the helix α1 and surrounding loops, while residues of ALK1 and V2 ALK2^Mimic-ALK1^ can form alike interactions to BMP9 (interaction site II & III) as shown by stick representation and distance measurements (using PyMOL).

To elucidate possible interactions of the ALK2 variants, we designed a homology model of each variant and docked it to BMP9 based on the ALK1/BMP9/ACVR2B complex (PDB:4FAO) and AlphaFold2 predictions of ALK2 (**Fig. S10-11, Fig.5E-F**)(Jumper et al., 2021; Townson et al., 2012; Varadi et al., 2022). Structure comparison underlined the incapability of the ALK2 helix α1 to form strong bonds at the wrist epitope (**Fig. 5E-F**). However, helix α1 of V2 but not of V1 ALK2^Mimic-ALK1^ can form interactions with BMP9, similarly as in ALK1 (**Fig. 5F, Fig. S10B**). Substitution of the HERR motif in V1 ALK2^Mimic-ALK1^ is likely insufficient due to the rigidity and/or altered conformation of surrounding residues (e.g., Thr_V1_179 *versus* Gly_V2/ALK1_79), prohibiting the conformations of the mutated residues needed for interacting with BMP9 (**Fig. S11B**). Although the binding interface of V3 ALK2^Mimic-ALK1^ and BMP9 was thought to become more like ALK1, the F2 loop differs in its conformation when comparing ALK1 and V3 ALK2^Mimic-ALK2^, possibly due to the ALK2-derived overall fold and surrounding residues **(Fig. S11C).**This likely explains the observed low binding of V3 ALK2^Mimic-ALK1^ to BMP9 (**Fig. 5C**). Together this indicates that exchanging critical residues of the helix α1 of ALK2 with ALK1 residues transfers BMP9 sensitivity.

## Discussion

Interrogating cell surface receptor biology in living cells, including their distribution, stoichiometries, ligand-binding efficacies and downstream activation has been pushed in the last years mainly by state-of-the-art microscopy techniques (Borgarelli et al., 2021; Kemter et al., 2021). In all these instances, the choice of labelling the protein of interest is of immense importance (Sahl, Hell, & Jakobs, 2017). While traditional fluorescent protein tagging is powerful, it comes with several drawbacks, such as delay in chromophore maturation, observation of intracellular signals, limited choice of different colors (especially in the far-red spectrum) and reduced performance in super-resolution nanoscopy due to bleaching. Small organic dyes overcome this issue, which are more robust and brighter, and can be selectively targeted in a temporal manner to self-labelling enzymes, such as SNAP- and Halo-tags. Spatial resolution can furthermore be achieved by using membrane-impermeable substrates and dyes that restrict labelling to extracellularly exposed tags. Even more, the possibility to use orthogonal tags allows the use of different colors, rapidly adding to the portfolio of multiplexed experiments to visualize receptor pairs and their ligand interaction in live cells. Recent examples include the determination of SNAP-mGluR4 co-localization (with Ca_v_2.1 and Bassoon) by two-color dSTORM microscopy (Siddig et al., 2020), the compositional diversity of differently tagged-kainate receptor assembly by three-color single molecule TIRF microscopy (Selvakumar et al., 2021), and the endocytosis and turnover of Halo-TfR by widefield imaging (Jonker et al., 2020). Alternatively, visualization of cell surface ligand binding, endo- and exocytosis, as well as receptor clustering was achieved using ligands covalently linked to fluorophores (Alborzinia et al., 2013; Ast et al., 2020; Nanda & Lorsch, 2014; Paarmann et al., 2016; Ramachandran et al., 2021; Trujillo et al., 2020). For this, ligands were either directly labeled at deprotonated primary amines using NHS-activated fluorophores (Alborzinia et al., 2013; Nanda & Lorsch, 2014), or through introduction of an N- or C-terminal cysteine residue allowing for site-directed maleimide coupling of the dyes (Ast et al., 2020; Paarmann et al., 2016).

Herein, we combined N-terminally Halo- and SNAP-tagged BMP and TGFβ receptors with fluorescently labeled growth factors that allowed us to measure and to visualize ligand binding to cell surface receptors in living cells. Whereas ligand receptor affinities are commonly assessed using *in vitro* methods such as surface plasmon resonance measurements (Patching, 2014), we attempted to measure ligand binding on living cells with the ability to resolve subcellular ligand binding events. Applying this *Ligand Surface Binding Assay* (LSBA) allowed for binding measurements under more physiological conditions with a focus on the plasma membrane. While in this study we focused on the binding properties of single receptors, future studies can include additional binding partners. Whereas crosslinking experiments using radio-labelled growth factors have been performed as one of the first methods to define high- and low-affinity BMP/TGFβ receptors affinities (Lin, Wang, Ng-Eaton, Weinberg, & Lodish, 1992; Mitchell & O’Connor-McCourt, 1991; Nishitoh et al., 1996; Rodriguez et al., 1995), the use of fluorescently labeled ligands increases the field of application to multiple microscopy techniques. By using labelled Activin A it was recently shown that ligand-mediated receptor clustering depends on the presence of ACVR2A/B (Ramachandran et al., 2021). Earlier studies using labelled BMP2 gave insights into ligand uptake and release (Alborzinia et al., 2013; Paarmann et al., 2016; Trujillo et al., 2020).

To test whether N-terminally Halo-/SNAP-tagged receptors are still capable of ligand binding we performed the LSBA using different members of the TGFβ ligand family (Activin A, TGFβ1 and BMP9), which are known to possess high affinities towards different type I or type II receptors. Our findings from LSBA measurements using Activin A-Cy5 are in good agreement with previous binding and crosslinking studies using radiolabeled ligands, highlighting that only the BMP/Activin type II receptors are capable of Activin A binding and they do not require the presence of type I receptors (Attisano et al., 1992; Donaldson et al., 1992; Rejon et al., 2013). SPR analysis confirmed that ACVR2B^ECD^ exhibited the highest affinity (K_D_ = 1.1 nM) followed by ACVR2A^ECD^ (K_D_ = 5.7nM) and BMPR2^ECD^ (K_D_ = 59 nM) (Heinecke et al., 2009). Receptor dimerization will further enhance Activin A binding as indicated by SPR studies using ACVR2A-Fc (K_D_ ^~^130 pM) and BMPR2-Fc (K_D_ ^~^14 nM)(Rejon et al., 2013), which was further confirmed by a comparison of Activin A binding to a single ECD ACVR2B-Fc (K_D_ = 108 pM) versus a homodimeric (ACVR2B)_2_-Fc (K_D_ = 12.5 pM)(Goebel et al., 2022). Interestingly, BMPR2-Activin A complexes have been suspected to be more transient due to their higher dissociation rate, compared to ACVR2A-Activin A (Rejon et al., 2013). These differences are confirmed by sequence comparison of the ACVR2B-Activin A interface with ACVR2A and BMPR2 ECD sequences. While ACVR2A shares mostly the same 2/3 loop with ACVR2B, which contains polar and charged amino acids that are critical for Activin A binding, BMPR2 has an alternative 2/3 loop (Thompson et al., 2003). We could show that substitution of these critical residues elevated Activin A binding to BMPR2 by ^~^3 fold. In the same light, substitution (H87A) of a sterically unfavorable Histidine located in the 2/3 loop led to a 1.5 fold increase of ^125^I-Activin A surface binding in CHO cells (Rejon et al., 2013). Together this highlights that the LSBA is suitable for mutant screening and assessment of high affinity ligand binders.

We next performed fluorescent labeling of BMP9 and TGFβ1, using a NHS-ester activated state-of-the-art silicon rhodamine derivate SiR-d12, which is characterized by high brightness and increased lifetime, suitable for confocal, life-time and super resolution microscopy (Roßmann et al., 2020). Both BMP9-SiR-d12 and TGFβ1-SiR-d12 bound their known high affinity receptor, i.e. ALK1 and TGFBR2, respectively (Cheifetz et al., 1987; David et al., 2007; Massague et al., 1992). Interestingly, whereas BMP9 is supposed to signal via ALK2 in non-endothelial cells that lack ALK1 expression (Kang et al., 2004; Luo et al., 2010; Olsen et al., 2014; Salmon et al., 2020), we did not observe BMP9 binding by any other type I receptor except ALK1. This is in line with the described difference of ALK1 being ^~^1800 times more sensitive to BMP9 than ALK2 as determined by SPR (Salmon et al., 2020). BMP9 is secreted into the blood stream from stellate cells in the liver (Miller, Harvey, Thies, & Olson, 2000). In the vasculature BMP9 circulates at concentrations (2-12 ng/mL) that are ^~^100 times higher than the EC_50_ of BMP9 to ALK1 (EC_50_ = 50 pg/mL, 2 pM)(David et al., 2008; David et al., 2007), highlighting that endothelial cells, which express predominantly ALK1 as type I receptor, are exposed to saturating BMP9 conditions. BMP9 signaling via ALK2 was reported to play a role in a variety of non-endothelial cells (Mostafa et al., 2019). Mechanistically, this was supported by radioactive crosslinking studies which observed BMP9-ALK2 binding in myoblasts and breast cancer cells (Scharpfenecker et al., 2007). Further, the authors highlighted that co-expression of BMP type II receptors in COS-7 cells enhances BMP9 binding to ALK2. While local BMP9 expression has been described in the developing murine CNS (Lopez-Coviella, Berse, Krauss, Thies, & Blusztajn, 2000), the predominant source of BMP9 in the human body is circulating BMP9 originating from the liver (David et al., 2008). Therefore, ALK2-dependent BMP9 signalling in non-endothelial cells could play a prominent role in the context of leaky or injured vessels accompanied by blood spillage into the tissue. In line with multiple reports of BMP9 influencing tumor progression (Mostafa et al., 2019), tumors are characterized of immature leaky blood vessels (Azzi, Hebda, & Gavard, 2013). Similarly, BMP9 could fulfil its previously described pro-osteogenic function (Lamplot et al., 2013) in the context of bone fracture healing.

Most of the ALK1-BMP9 interface is unique (Townson et al., 2012), which is why we sequentially substituted key residues of ALK2 with corresponding residues of ALK1. Particularly substituting the amino acids of the pre-helix loop and the helix α1 were sufficient to increase BMP9 binding to V2 ALK2^Mimic-ALK1^. Interestingly, we did not detect ACVR2B dependent BMP9 binding, while SPR studies using ACVR2B-Fc and ALK1-Fc indicated a similar affinity of BMP9 for both receptors (Townson et al., 2012). A potential explanation could be that a stable dimer of ACVR2B, as present in the Fc-fusion protein, is required for high affinity binding of BMP9, which might not be the case for the full-length receptors at the plasma membrane. While a comparative study for ACVR2B binding to BMP9 has to our knowledge not been performed, monomeric ALK1-ECD (K_D_ ^~^ 71.6 pM) possesses a lower affinity to BMP9 compared to dimerized ALK1-Fc (K_D_ ^~^ 48.1 pM) (Salmon et al., 2020). It is unlikely that BMP9 binding to ACVR2B was negatively influenced by the N-terminal HALO-tag, as the same receptor was able to bind Activin A normally (Salmon et al., 2020; Thompson et al., 2003; Townson et al., 2012). However, an independent SPR study highlighted that the affinity of Activin A (K_D_ ^~^ 9.2 pM) towards ACVR2B-Fc is eleven times higher than the affinity of BMP9 (K_D_ ^~^ 116.1 pM) (Li et al., 2021). Furthermore, administration of ACVR2B-Fc as a GDF8/Myostatin inhibitor (ACE-031) in a clinical trial to treat Duchenne muscular dystrophy had to be stopped due to the occurrence of telangiectasias (Campbell et al., 2017), typical of deficient BMP9-signaling in the vasculature (Cunha, Magnusson, Dejana, & Lampugnani, 2017). Together this highlights, that BMP9 efficiently binds to monomeric ALK1 at low concentrations, as well as to hetero- (e.g. ALK2/ACVR2B) or homodimeric receptor complexes (ACVR2B/ACVR2B).

In summary, we established an imaging platform that allows us to revise and test ligand receptor affinities in a cellular context. Due to the nature of the self-labeling Halo- and SNAP-tag, BMP and TGFβ receptors can now be visualized at a stoichiometric ratio using custom dyes, which meet the requirements for the respective experiments, e.g impermeability for surface-only stainings, increased photo-stability or STEDability (Birke et al., 2022; Roßmann et al., 2020). Future studies can now address (1) differences in ligand-receptor binding of monomeric receptors vs. receptor complexes, (2) binding behavior of other TGFβ-family ligands using the incorporation of unnatural amino acids at the C-terminus allowing site-directed coupling and positioning of fluorophores, (3) visualization of endogenous receptor populations in cells and in tissues using fluorescently labeled high affinity ligands as replacement for insufficient antibodies.

## Material and Methods

### Restriction cloning

First hBMPR2-LF-SNAP/Halo and mUnc5b-SNAP/Halo were generated by substitution of meGFP in hBMPR2-LF-meGFP and mUnc5b-meGFP *(doctoral thesis J. Jatzlau)* with SNAP or Halo using Sec-61-Halo and pSNAPf as a template (Bottanelli et al., 2016). For the generation of the N-terminally Halo- and SNAP-tagged BMP / TGFβ receptor library, hALK1-hALK6, rALK7 and hALK2-R206H were subcloned into pcDNA3.1 mUnc5b-SNAP/Halo and hACVR2A, hACVR2B, hTGFBR2 were subcloned into pcDNA3.1 hBMPR2-LF-SNAP/Halo in between EcoRI and NotI thereby replacing the respective receptor ORF. All cloning PCRs were carried out on a Peltier Thermal Cycler PTC-200 and the primers used are listed in (Appendix Tab. 1). The elongation time was adapted according to the product size (Phusion Pol. = 1 kb/min). PCR products were resolved by agarose gel electrophoresis and purified using NucleoSpin Gel and PCR Clean-up kit, according to the manufacture’s guidelines. For restriction cloning, PCR products and 3 μg of the destination vector were digested overnight with 1 μL (^~^ 20 Units) of each restriction enzyme. Successful restriction digest was validated on an agarose gel, from where cleaved products were purified. Subsequently, 50 ng destination vector and 300 ng insert were ligated for 10 min at room temperature with T4 ligase (^~^ 40 Units) in 10 μL H_2_O. After 5 min heat inactivation at 65°C, the total volume of ligation mix was added to 50 μL chemically competent DH5α or TOP10 *E.coli* and incubated for 30 min on ice. After a 90 seconds heat shock in a 42°C water bath, the bacteria were put back on ice for 2 min. Next, the bacteria were shaken for 1 hour at 37°C and 180 rpm in 1 mL LB medium w/o antibiotics. Finally, bacteria were plated on LB-agar plates with antibiotics and grown overnight at 37°C. On the next day, 3 mL LB medium with the respective antibiotics were inoculated with bacterial colonies and grown at 37°C and 180 rpm overnight, followed by plasmid DNA purification and Sanger sequencing.

### Cell culture

COS-7 cells were obtained from the German Collection of Microorganisms and Cell Cultures (DSMZ) and cultured in Dulbecco’s Modified Eagle’s Medium (DMEM) supplemented with 10% FCS, 2 mM L-glutamine and penicillin (100 units/mL) / streptomycin (100 μg/mL) (DMEM full medium) in a humidified atmosphere at 37 °C and 5% CO_2_ (v/v). COS-7 cells were maintained in T175 flasks and cells were split 1:3 or 1:5, depending on need and were kept sub-confluent. For passaging, cells were washed once with PBS before being removed from the flasks surface with trypsin/EDTA (0.05/0.02% in PBS).

### Transient transfection with expression plasmids

For microscopy studies of COS-7 cells, cells were transfected with Lipofectamine 2000 according to the manufacturer’s instructions. 200,000 cells / 12-well were seeded in 1 mL DMEM full medium. On the following day, cells in each well were transfected with a total amount of 500 ng DNA. Subsequent experimental procedure took place 24 hours post transfection, unless indicated otherwise.

### SDS-PAGE & Western-blotting

For sodium dodecyl sulfate polyacrylamide gel-electrophoresis (SDS-PAGE), treated cells were lysed in 150 μL Laemmli buffer (Laemmli, 1970) and frozen at −20 °C. The lysate was pulled through an 18-gauge syringe and boiled for 10 min at 95°C. 10% polyacrylamide gels were cast in advance and stored at 4 °C until usage. Separated by their molecular weight, proteins were transferred onto methanol-activated PVDF membranes by Western-blot. Membranes were blocked for 1 hour in 0.1% TBS-T containing 3% w/v BSA, washed three times in 0.1% TBS-T and incubated with indicated primary antibodies overnight at 4°C. Primary antibodies: anti-GAPDH (Cell Signaling; #2118; monoclonal rabbit antibody), anti-Halo (ProMega; #G9211; monoclonal mouse antibody) anti-SNAP (Invitrogen; #CAB4255; polyclonal rabbit antibody) were applied at a 1:1000 dilution in 3% w/v bovine serum albumin (BSA)/fraction V in TBST. For HRP-based detection, goat-α-mouse or goat-α-rabbit IgG HRP conjugates (± 0.8 mg/ml, Dianova; #111-035-144, #115-035-068) were used at a dilution of 1:10,000. Chemiluminescent reactions were processed using WesternBright Quantum HRP substrate (advansta) and documented on a FUSION FX7 digital imaging system.

### Halo- & SNAP-tagged receptor staining & ligand visualization

For visualization of COS-7 cells transiently expressing Halo- and SNAP-receptor fusion proteins, 200,000 cells were transfected on glass cover slips (confocal: 18 mm ø or STED: precision #1.5H 10 mm ø) with desired constructs. For live cell imaging, 200,000 COS-7 cells were seeded on glass-bottom culture dishes (35 mm ø). 24 hours post-transfection, cells were washed once with PBS and incubated, depending on the experimental setup, with a fluorescent SNAP- and / or a fluorescent Halo-ligand in the absence or presence of fluorescent growth factors for 30 minutes. The following commercially available cell-impermeable fluorescent ligands were utilized: HaloTag Fluorescent Ligand Alexa Fluor 488 (Promega, #G1002) in the following abbreviated as CA-Alexa488; SNAP-Surface 594 (NEB, #S9134S) in the following abbreviated as BG-ATTO594 and SNAP-Cell 647-SiR (NEB, # S9102S), in the following abbreviated as BG-SiR. Surface receptor staining was carried out at 4 °C, whereas incubation at 37 °C took place, when studying growth factor binding on living cells. Cells were washed once with PBS, before fixation with 100% methanol for 5 minutes. Cells were washed with PBS once more and mounted with Fluoromount G (confocal microscopy).

### Preparation of fluorescently labelled recombinant Activin A

Mature Activin A was prepared by refolding from inclusion bodies as described before for Atto-647 and CF640R-labelled activin A (Ramachandran et al., 2021) with the exception that mature activin A was untagged and the label used here was NHS-activated Cy5 (Lumiprobe, cat. no 13020). The protein:dye ratio was 1:2 in the labelling reaction with 76 μM Activin A, 153 μM NHS-Cy5, 50 mM HEPES pH 7.4, and 30% acetonitrile. Labelling was for 3 hours at 23 °C and labelled protein was purified by reversed phase chromatography, fractions with labelled protein divided into 5 μg aliquots and vacuum dried.

### Cell stimulation with growth factors

rhBMP9/GDF2 and rhTGFβ1 (PeproTech, Hamburg, Germany) were reconstituted and stored according to manufacturer’s instructions. For microscopical binding studies, COS-7 cells were stimulated with 2 nM Activin A-Cy5. For microscopical binding studies (COS-7) of SiR-d12- or labeled growth factors, cells were stimulated with 0.3 nM TGFβ1-SiR-d12 and 0.3 nM BMP9-SiR-d12 if not indicated otherwise.

### Fluorescent growth factor labeling & Ligand Surface Binding Assay (LSBA)

For fluorescent labeling of BMP9 and TGFβ1, *N*-hydroxysuccinimidyl deuterated silicon rhodamine (NHS-SiR-d12) (synthesis see Supporting Information) was taken up in DMSO to a final concentration of 1 mM. Lyophilized rhBMP9 (Peprotech) and rhTGFβ1 (Peprotech) or were reconstituted in 0.2 M Sodium bicarbonate (reaction buffer) to obtain a final concentration of 2 μM. NHS-SiR-d12 was then added to the reconstituted ligands at a 5-fold molar excess. Final concentrations being at 2 μM (Protein) and 10 μM (NHS-Dye). The mixture was allowed to incubate at room temperature for 4 h. Meanwhile, a 0.5 mL Amicon Ultra 10 kDa molecular weight cutoff column (Merck Millipore, UFC501024) was calibrated with reaction buffer and centrifuged at 14,000 × *g* at 4°C without drying the column. The reaction mixture was then gently applied onto the pre-calibrated column and centrifuged for 4 minutes, while never allowing the column to dry. Subsequently, the column was gently washed with reaction buffer *ad* 500 μL and centrifuged as described before. This washing procedure was repeated twice. Afterwards, 50 μL sterile Millipore H_2_O was added onto the column, before the column was inverted and put in a fresh elution tube and centrifuged as described above. The column was rinsed with 30 μL sterile Millipore H_2_O and eluted as in the previous step. Protein concentration was determined with a Nano Drop 2000 spectrophotometer (ThermoFischer). For visualization of SiR-d12-ligand binding on COS-7 cells, cells were transfected as described above with corresponding high- and low-affinity Halo-receptors. The day after, cells were washed with PBS and, additionally to fluorescent Halo-ligand CA-Alexa488, simultaneously incubated with fluorescent growth factors at previously indicated concentrations for 30 minutes at 4 °C. Subsequently, cells were washed once with ice-cold PBS before fixation with 100% methanol for 5 minutes. Cells were washed again with PBS and mounted with Fluoromount G (confocal microscopy) or ProLong Gold Antifade Mounting reagent (STED microscopy).

### Confocal & STED microscopy

Confocal and STED data of fixed or living COS-7 cells were acquired with the Expert Line STED Microscope from Abberior. Confocal images of COS-7 cells expressing Halo-tagged receptors stained with CA-Alexa488 and fluorescent ligands (Cy5- & SiR-d12-labeled) were acquired using 485 nm (20% laser power) and 640 nm excitation (20% laser power). STED images were acquired using 561 nm excitation for ATTO 594 (20% laser power) and 640 nm excitation for SiR (2% laser power). A 775 nm STED laser at 10% laser power was used to deplete both dyes. Live cell confocal imaging of COS-7 cells transiently expressing ACVR2B-Halo was carried at a Abberior Expert Line STED microscope at physiological temperature of 37 °C using a stage-top incubator (okolab). Cells were stained with 0.5 μM Halo-tag substrate CA-AlexaFluor488 (Promega) for 30 minutes at 37°C, washed extensively and kept on ice in HEPES buffer pH 7.4 until imaging. Focussing on transfected and stained cells, Activin A-Cy5 was added to the cells at 2 nM and dual-color images were acquired for two minutes every two seconds. Confocal and STED Images were acquired with the following settings in the Abberior Imspector Acquisition & Analysis Software v16.3: objective lens: 100X NA1.4 (oil) [UPLS], pinhole: 1.0 AU, range: 75 μm × 75 μm, confocal pixel size: 60 nm, STED pixel size: 20 nm, pixel dwell time: 5/10 μs.

### Image analysis & semi-automated quantification with Fiji ImageJ

Confocal raw data were post-processed and adjusted for color and contrast (linear adjustments maintained for confocal datasets represented within one figure) using Fiji (ImageJ) software and Adobe Photoshop (Adobe Systems). Surface binding quantification of Activin A-Cy5, BMP9-SiR-d12 and TGFβ1-SiR-d12 on COS-7 cells transiently expressing SNAP- & Halo-Receptor constructs was performed with Fiji. Per cell, four regions of interest (ROI) (100 μm^2^) were chosen and the raw integrated intensity (RawIntDen) of each ROI was measured both in receptor and ligand channels. Per condition, 10-30 cells were quantified in 3 independent experiments. Receptor-ligand binding was calculated relative to intensity values of untransfected COS-7 cells, representing endogenous ligand-receptor binding. Linear regression and correlation analysis of RawIntDen values of receptors and ligands was performed and plotted in GraphPad Prism 8.0 (GraphPad Software Inc.). Maximum RawIntDen values of receptors were normalized to 1. All used scripts are available under https://github.com/Habacef/Supplementary-scripts-for-LSBA.

### Statistical analysis

All statistical tests were performed using GraphPad Prism version 9.3 software and are listed in the figure legends. Normal distribution of data sets were tested with the Shapiro-Wilk normality test. In cases of failure to reject the null hypothesis, the ANOVA and Tukey’s post hoc test were used to check for statistical significance under the normality assumption. For all experiments statistical significance was assigned, with an alpha-level of p < 0.05.

### Homology modeling

The receptor variant BMPR2^Mimic-ACVR2B^ was modeled starting from a BMPR2 x-ray structure (PDB: 2HLQ, (Mace et al., 2006)). The receptor variants V1 ALK2^Mimic-ALK1^, V2 ALK2^Mimic-ALK1^ and V3 ALK2^Mimic-ALK1^ were modeled starting from the AlphaFold2 prediction of ALK2 (Jumper et al., 2021; Varadi et al., 2022), as the topology of the ECD of ALK2 is not yet experimentally solved. The starting structures were prepared using a coordinate constrained relaxation protocol to obtain optimized H-bonds based on the ref2015_cst score function within RosettaScripts (Fleishman et al., 2011). The GreedyOptMutationMover (King et al., 2014; Nivón, Bjelic, King, & Baker, 2014) was used to generate 500 replicas of each variant via insertion of mutation (and if applicable a short, up to 3 AA long, loop modeling). Within the same RosettaScript, a full FastRelax (Khatib et al., 2011; Maguire et al., 2021) with the filters “BuriedUnsatHBonds” and “PackStat” was performed for each generated variant. The ten best variants were discriminated by plotting the RMSD of all heavy backbone atoms vs. the Rosetta-Score (using scoring function ref2015) of the final 500 variants. The RMSD was calculated against the input structures (either BMPR2 or ALK2) using the RMSDMetric mover (Adolf-Bryfogle et al., 2021) in RosettaScripts. The ten variants with low REU-values (score) and low RMSD were visually inspected, and the most reliable structure based on convincing intramolecular interactions and comparison to known BMP-receptor-ECDs was used for receptor-ligand-docking. The described plots together with cartoon representations of all variants are shown in Fig. S8A and Fig. S10A. All used RosettaScripts can be found in an open-source repository under https://github.com/Habacef/Supplementary-scripts-for-LSBA.

### Receptor ligand docking

All homology models of receptor variants, but also the X-ray structure of BMPR2 and the AlphaFold prediction of ALK2 were docked to Activin A and BMP9, respectively. BMPR2 and BMPR2^Mimic-ACVR2B^ were superimposed to Activin A-bound ACVR2B (X-ray structure PDB: 1S4Y). ALK2, V1 ALK2^Mimic-ALK1^, V2 ALK2^Mimic-ALK1^ and V3 ALK2^Mimic-ALK1^ were superimposed to BMP9-bound ALK1 (X-ray-structure PDB: 4FAO, (Townson et al., 2012)). As the receptor binding interfaces (Fig. S11A) are known, local Monte-Carlo-based protein-protein docking could be performed in 500 replicas. Here, first, the “Docking” mover was used with low resolution (backbone plus centroid) flags before a high resolution (full atom) docking (using ref2015 scoring function). Other parameters as distance perturbation and angle perturbation were left on default. The Interface RMSD (IRMSD) was plotted against the score of the interface residues (ISC). Here too, the best 10 hits showing the lowest IRMSD and lowest ISC were screened for the final docked homology model. The described plots together with cartoon representations of all variants are shown in Fig. S8B and Fig. S10B. All used scripts are available under https://github.com/Habacef/Supplementary-scripts-for-LSBA.

### Graphical schemes and figures

Figures were created with BioRender.com and *PyMOL*, Available at: http://www.pymol.org/pymol.

(Schrödinger, L. & DeLano, W., 2020.)

For additional information regarding materials and methods, see the Appendix Supplementary Methods.

## Acknowledgments

J.J. was supported by the Deutsche Forschungsgemeinschaft DFG (BSRT, SFB958) and the Einstein Center ECRT. P.K. acknowledges the support from Deutsche Forschungsgemeinschaft DFG (FOR2165; SFB1444), the Einstein Center ECRT, Morbus Osler Society and BMBF (PrevOP-Overload). We would like to acknowledge the assistance of the Core Facility BioSupraMol FU Berlin supported by the DFG. S.A.-S. was supported by the Excellence cluster REBIRTH, SFB958, by Deutsche Forschungsgemeinschaft (DFG) projects SE2016/7-2 and SE2016/10-1 and by the DZHK. F.B. was supported by the Deutsche Forschungsgemeinschaft DFG (SFB958).

## Author Contributions

J.J. and P.K. designed the study; J.J., J.B. and P.K. designed the labelling strategies and evaluated the data; W.B., J.J. and M.T. performed the experiments; L.O. performed receptor homology modeling & docking calculations; K.Ro. and J.B. produced NHS-SiR-d12; K.Ra. and M.H. produced the Cy5-labelled Activin A; F.B. provided expertise in STED microscopy; J.J., W.B. and P.K. wrote the manuscript; all authors commented on the manuscript.

## Data availability

The data supporting the findings of this study are available from the corresponding author upon request.

## Supplemental Figures

**Supplementary Figure 1:**
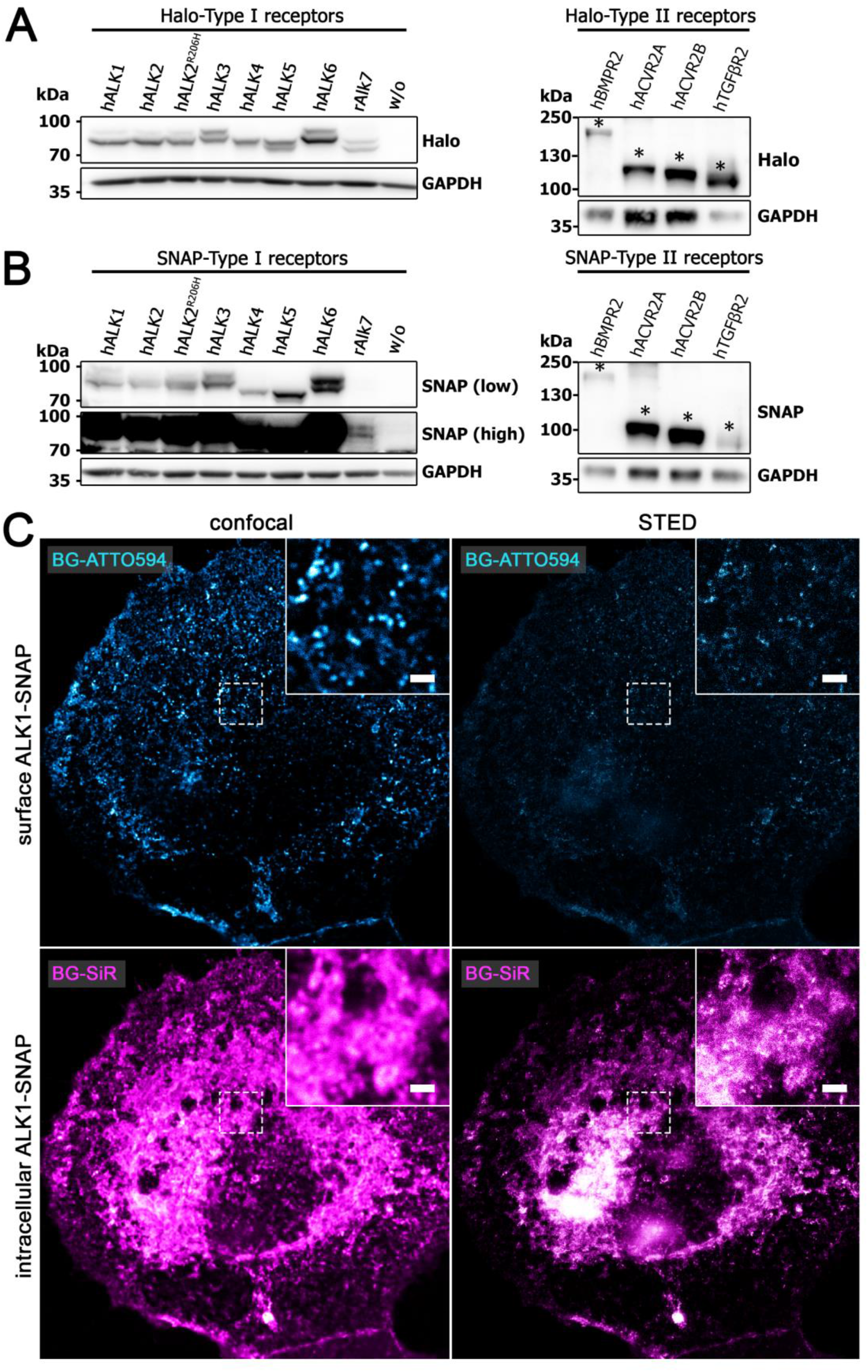
*Supplement to Fig.1*, **(A-B)** Expression of **(A)** Halo- and **(B)** SNAP-tagged receptors in COS-7 cells validated by Western blotting with specific antibodies directed against the Halo- or SNAP-tag, using GAPDH as loading control. **(C)** Discrimination between surface and cytosolic receptor populations. COS-7 cells transiently expressing ALK1-SNAP were 24 hours post transfection incubated with BG-ATTO594 (impermeable; cyan) followed by BG-SiR (permeable; magenta) incubation allowing for staining of the surface receptor population and the cytosolic receptor population, respectively. Confocal images (left) at a resolution of 60 × 60 nm/pixel compared with STED images (right) at a resolution of 20 × 20 nm/pixel. Scale bar ≙1μm.

**Supplementary Figure 2:**
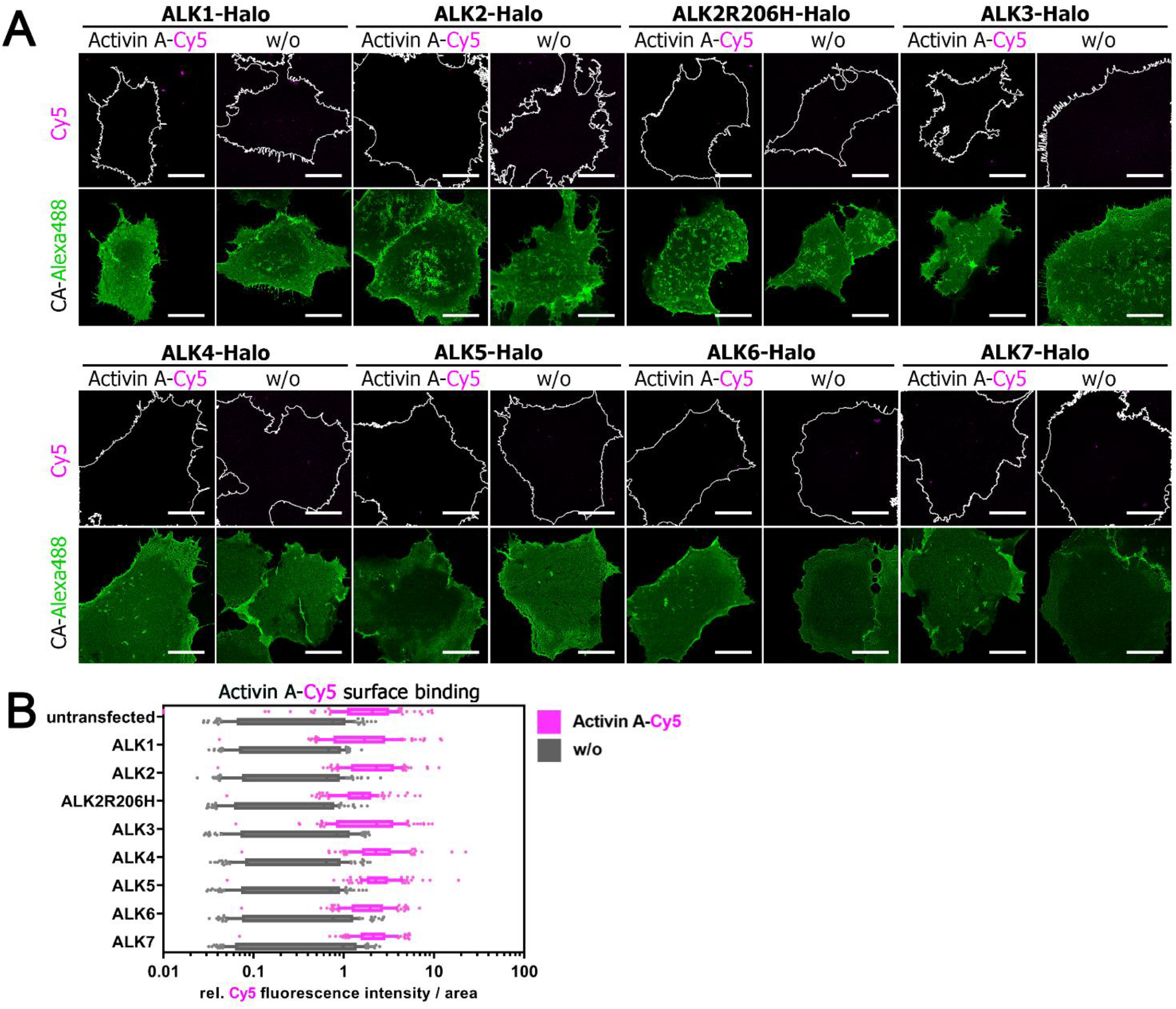
*Supplement to Fig.2*, COS-7 cells were seeded on coverslips and transfected with indicated Halo-tagged type I receptor constructs. 24 hours post transfection, cells were incubated with fluorescent Halo-tag substrate CA-Alexa488 (green) and Activin-Cy5 (magenta) for 30 minutes at 4 °C, fixated with methanol for 5 minutes at room temperature and mounted on glass slides. Cells were imaged at a confocal microscope and 10 cells per condition and replicate were analyzed with a semi-automated Fiji ImageJ macro pipeline for assessment of fluorescent growth factor binding (Activin A-Cy5 signal intensity) and fluorescence intensity of receptors (CA-Alexa488). Four ROIs of 100 μm^2^ were quantified in each cell. **(A)** Representative confocal microscopy images of COS-7 cells transiently expressing the Halo-tagged type I receptor library incubated with CA-Alexa488 and simultaneously stimulated with Activin A-Cy5 or PBS as control. Scale bar ≙ 20 μm. **(B)** Activin A-Cy5 surface binding represented as relative fluorescence intensity per area. Data is shown as F.I. ± SD. (n = 3 independent experiments) (CA: chloroalkane).

**Supplementary Figure 3:**
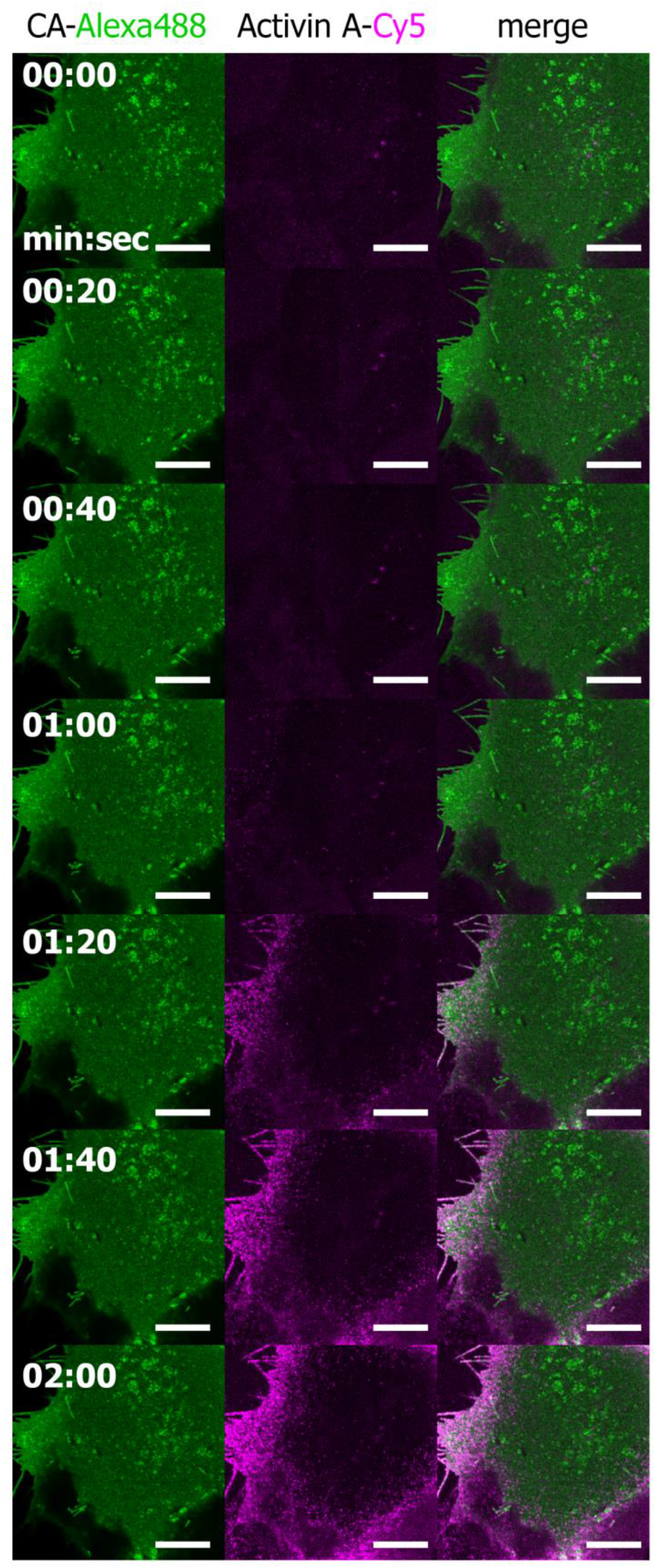
*Supplement to Fig.2*, Diffusion and binding of Activin A-Cy5 on COS-7 cells in live cell imaging (LCI). COS-7 cells were seeded on 35mm glass bottom dishes (Cellvis) and transiently transfected with ACVR2B-Halo. 24 hours post transfection, cells were stained with CA-Alexa488 for 30 minutes at 4 °C and stored on ice until image acquisition. Stimulation with 2 nM Activin A-Cy5 and imaging was performed in a LCI chamber (okolab) at 37°C. A total of 60 images were acquired over a time of 2 minutes.

**Supplementary Figure 4:**
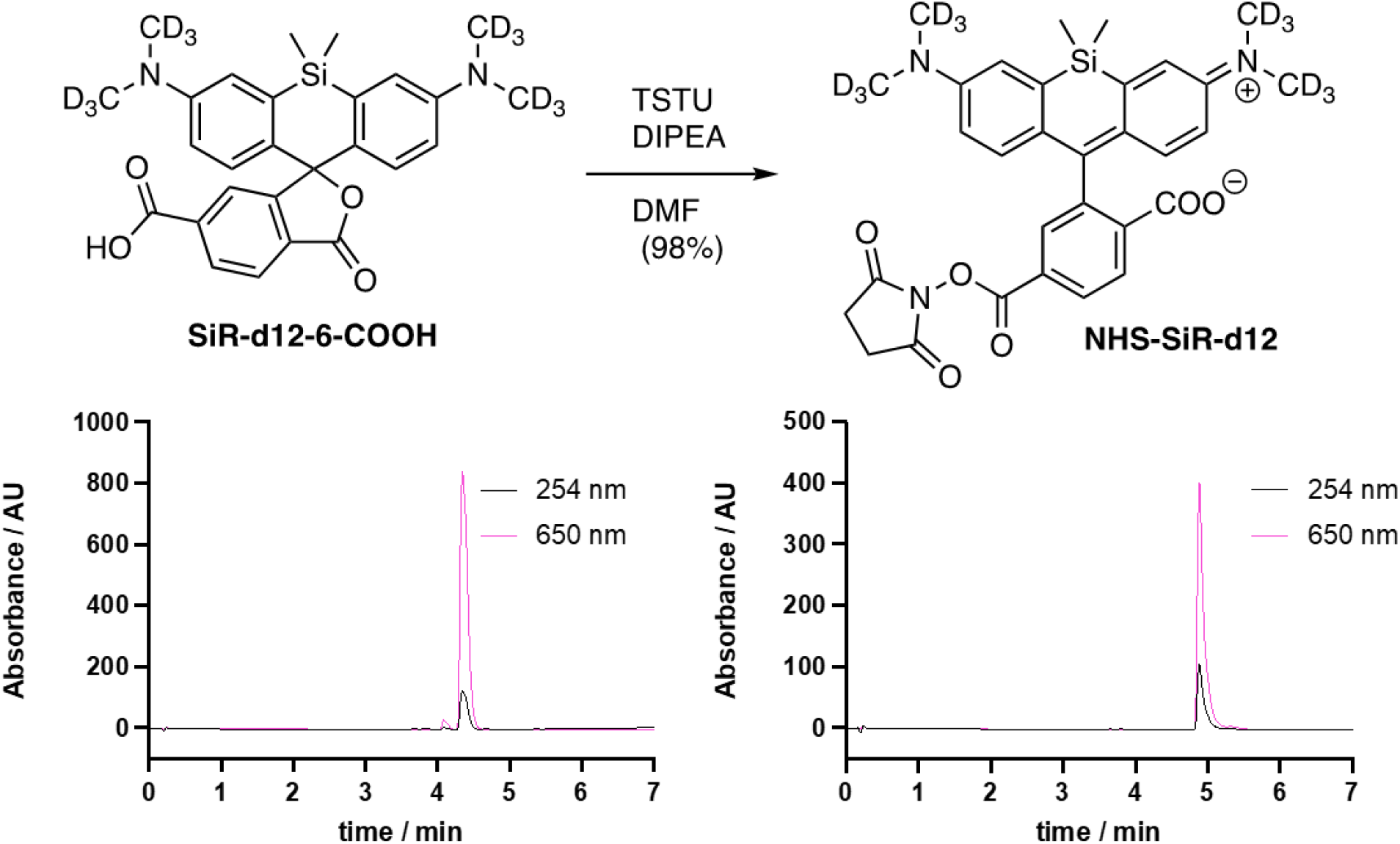
*Supplement to Fig.3*, Synthesis of NHS-SiR-d12. SiR-d12-6-COOH is activated by means of TSTU in DIPEA and DMF and the corresponding active ester is collected by RP-HPLC. LCMS of both substances below indicates purity.

**Supplementary Figure 5:**
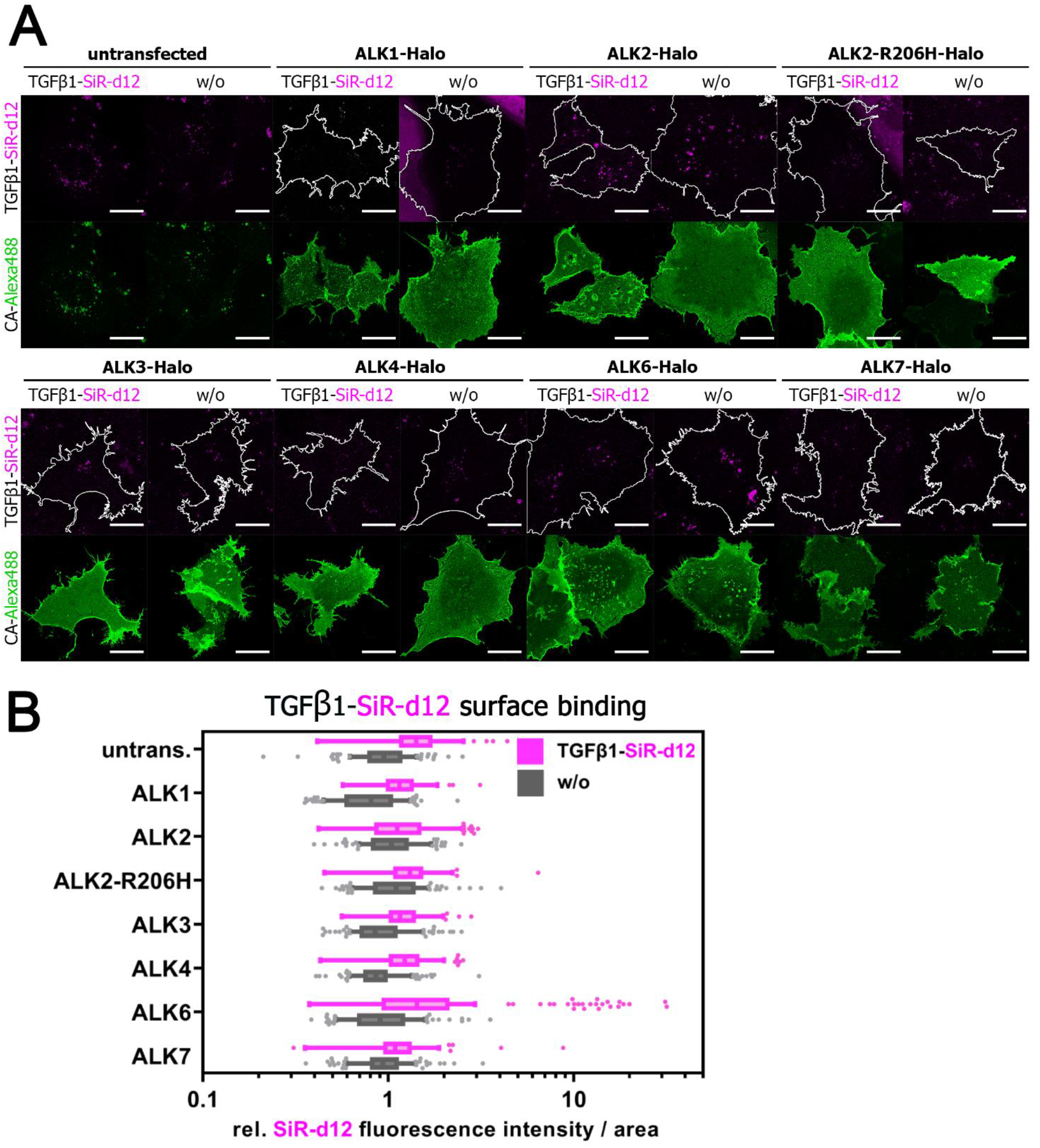
*Supplement to Fig.3*, Transiently transfected COS-7 cells expressing Halo-tagged type I receptor library were 24 hours post transfection simultaneously incubated with Halo-tag substrate CA-Alexa488 (green) and TGFβ1-SiR-d12 (magenta). **(A)** Representative confocal microscopy images. Scale bar ≙ 20 μm. **(B)** TGFβ1-SiR-d12 surface binding represented as relative fluorescence intensity per area. Data is shown as F.I. ± SD. (n = 3 independent experiments).

**Supplementary Figure 6:**
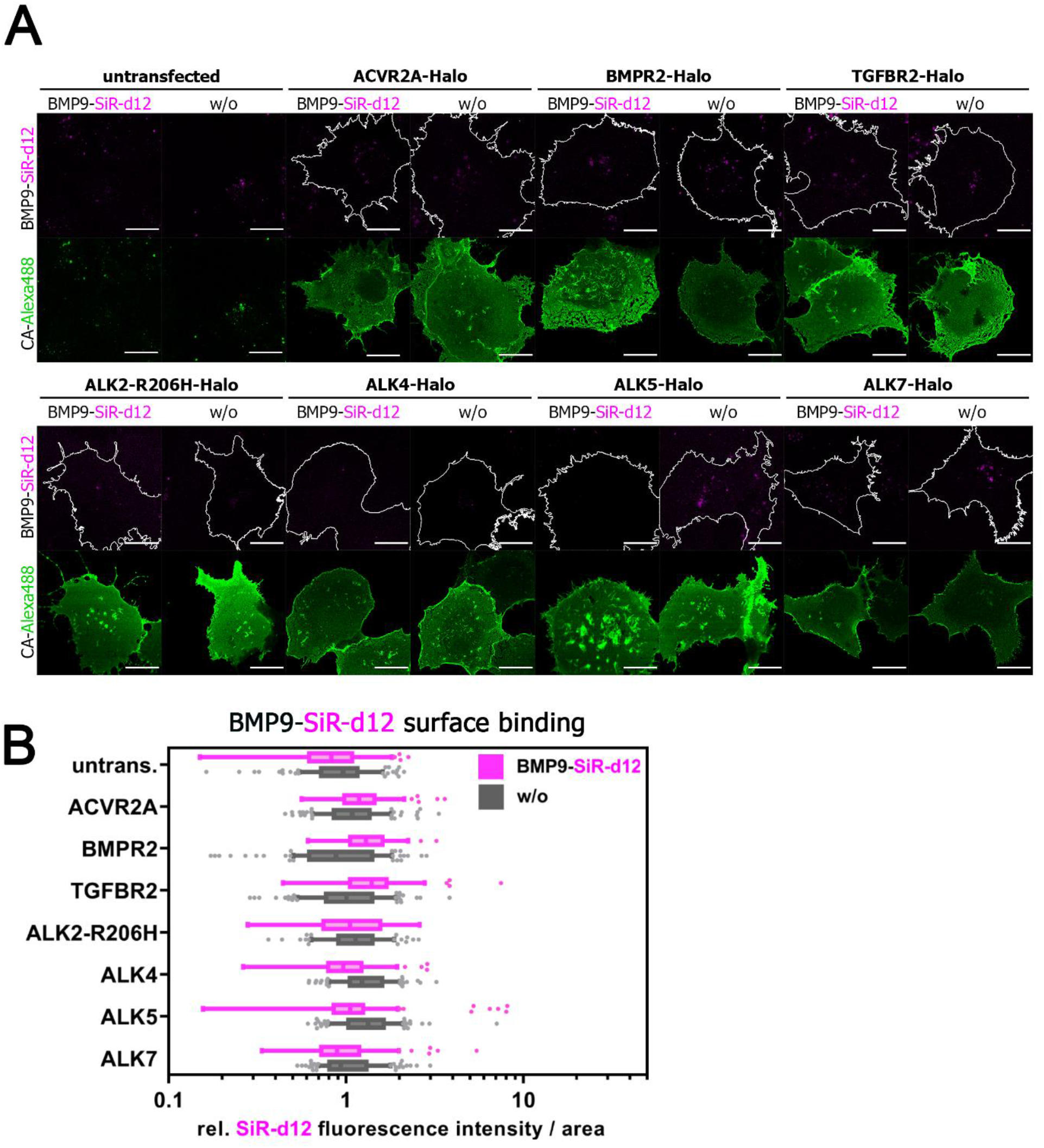
*Supplement to Fig.3*, Transiently transfected COS-7 cells transiently expressing Halo-tagged type II receptors ACVR2A-Halo, ACVR2B-Halo, BMPR2-Halo or type I receptors ALK2-R206H-Halo, ALK4-Halo, ALK5-Halo, ALK7-Halo were 24 hours post transfection simultaneously incubated with Halo-tag substrate CA-Alexa488 (green) and BMP9-SiR-d12 (magenta). **(A)** Representative confocal microscopy images. Scale bar ≙ 20 μm. **(B)** BMP9-SiR-d12 surface binding represented as relative fluorescence intensity per area. Data is shown as F.I. ± SD. (n = 3 independent experiments).

**Supplementary Figure 7:**
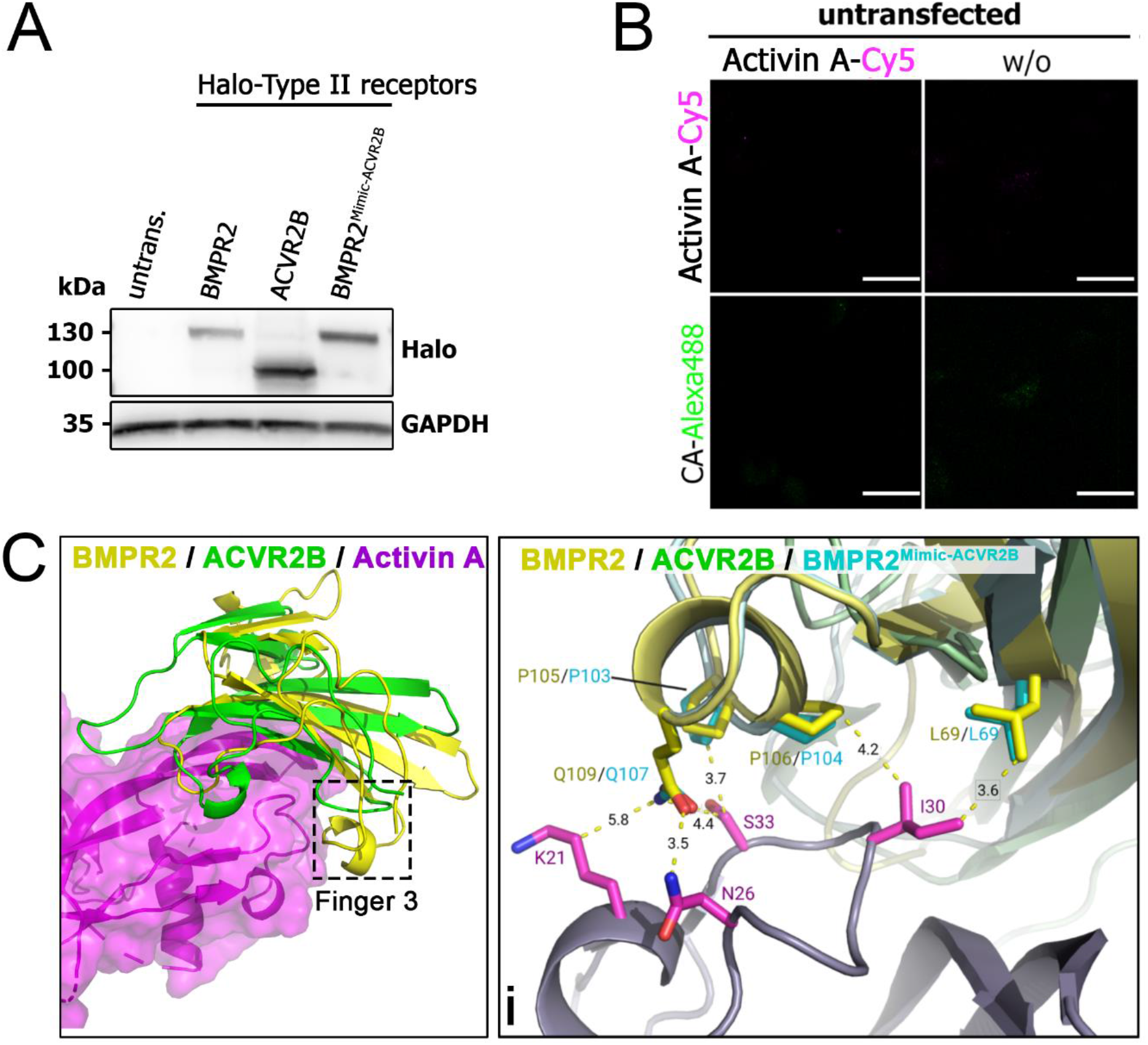
*Supplement to Fig.4* **(A)** Expression of Halo-tagged receptors BMPR2-Halo, ACVR2B-Halo and BMPR2^Mimic-ACVR2B^-Halo validated by Western blotting. Halo-tagged receptors transiently expressed in COS-7 cells validated by Western blotting with specific antibodies directed against Halo-tag, using GAPDH as loading control. **(B)** Untransfected COS-7 cells were 24 hours post transfection simultaneously incubated with Halo-tag substrate CA-Alexa488 (green) and Activin A-Cy5 (magenta). Representative confocal microscopy images of untransfected COS-7 cells incubated with CA-Alexa488 and simultaneously stimulated with Activin A-Cy5 or PBS as control. Scale bar ≙ 20 μm. **(C)** Overview (left) of BMPR2 (2HLQ) and ACVR2B in complex with Activin A (1S4Y) highlighting Finger 3 of BMPR2. Distance measurement within a stick and cartoon representation of the interface between Activin A at Finger1&2 and BMPR2 / BMPR2^Mimic-ACVR2B^ / ACVR2B at Finger 3 reveals a potential additional binding site present in both BMPR2 and BMPR2^Mimic-ACVR2B^, but not ACVR2B.

**Supplementary Figure 8:**
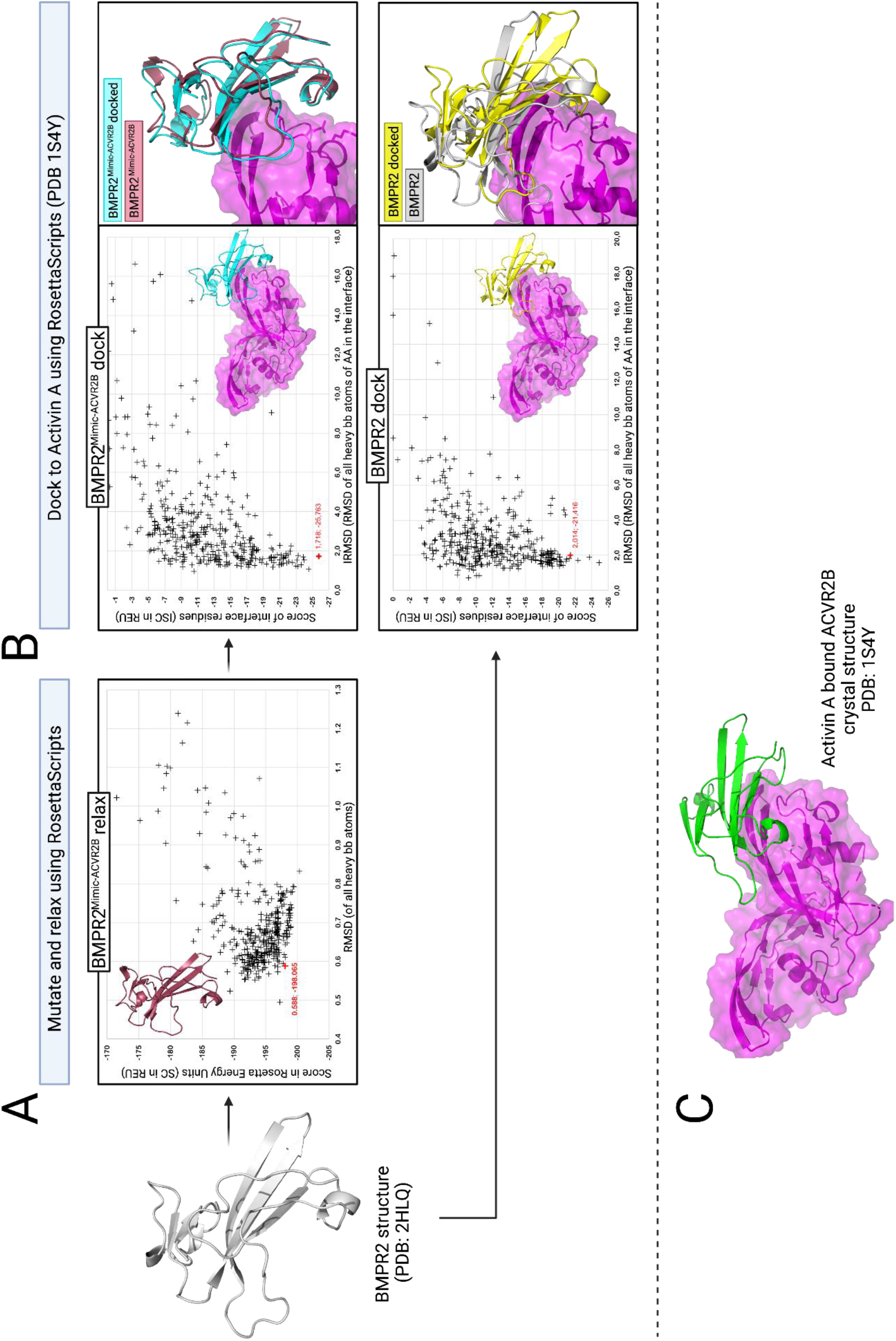
*Supplement to Fig.4* Simplified, schematic overview of the *in silico* homology modeling of BMPR2^Mimic-ACVR2B^ and docking of BMPR2 and BMPR2^Mimic-ACVR2B^ to Activin A using the Rosetta Commons Modelling Suite. **(A)** RMSD of all heavy backbone atoms vs. the Rosetta-Score (using scoring function ref2015) is plotted for 500 homology modelling attempts of BMPR2^Mimic-ACVR2B^ (derived from the BMPR2 crystal structure (2HLQ) by mutating and relaxing using the Rosetta Commons Modelling suite). The Datapoint of the final homology model (cartoon representation upper left corner of the plot) is highlighted in red. **(B)** After superimposing BMPR2 and BMPR2^Mimic-ACVR2B^ to Activin A bound ACVR2B (PDB: 1S4Y) and docking, the Interface RMSD (IRMSD) was plotted against the score of the residues (ISC) in the docking interface. Datapoints of the final dockings (surface & cartoon representation, lower right corner of the plot) are highlighted in red. A structural comparison of the final docking vs. a simple superimposition is shown right next to the plot for both BMPR2 and BMPR2^Mimic-ACVR2B^. **(C)** Cartoon and surface representation of the crystal structure 1S4Y of ACVR2B bound to Activin A.

**Supplementary Figure 9:**
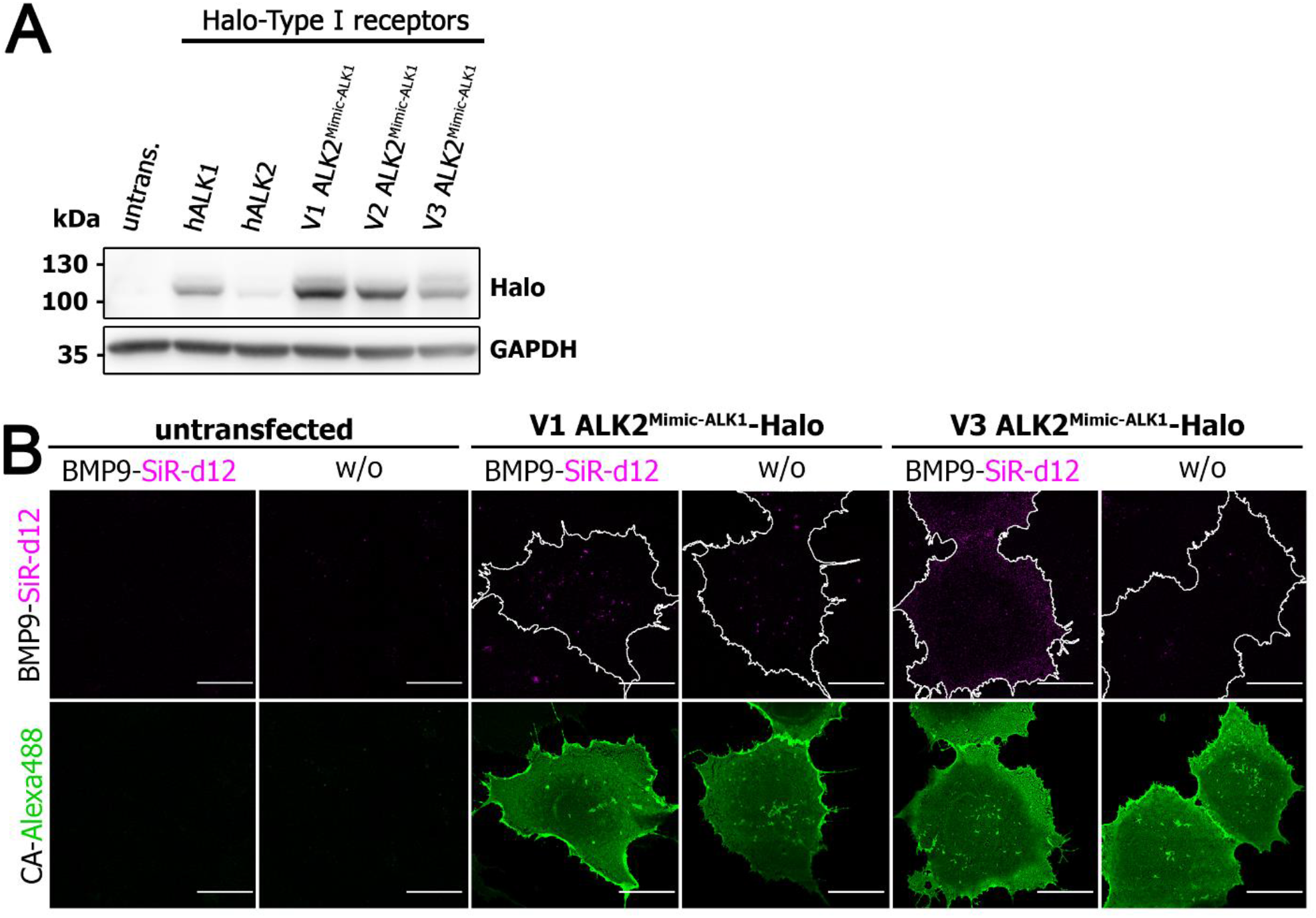
*Supplement to Fig.5* **(A)** Expression of Halo-tagged receptors ALK1-Halo, ALK2-Halo, V1 ALK2^Mimic-ALK1^-Halo, V2 ALK2^Mimic-ALK1^ and V3 ALK2^Mimic-ALK1^-Halo validated by Western blotting with specific antibodies directed against Halo-tag, using GAPDH as loading control. **(B)** Untransfected COS-7 cells or cells transiently expressing V1 ALK2^Mimic-ALK1^-Halo or V2 ALK2^Mimic-ALK1^-Halo were 24 hours post transfection simultaneously incubated with Halo-tag substrate CA-Alexa488 (green) and Activin A-Cy5 (magenta). Representative confocal microscopy images of untransfected COS-7 cells incubated with CA-Alexa488 and simultaneously stimulated with Activin A-Cy5 or PBS as control. Scale bar ≙ 20 μm.

**Supplementary Figure 10:**
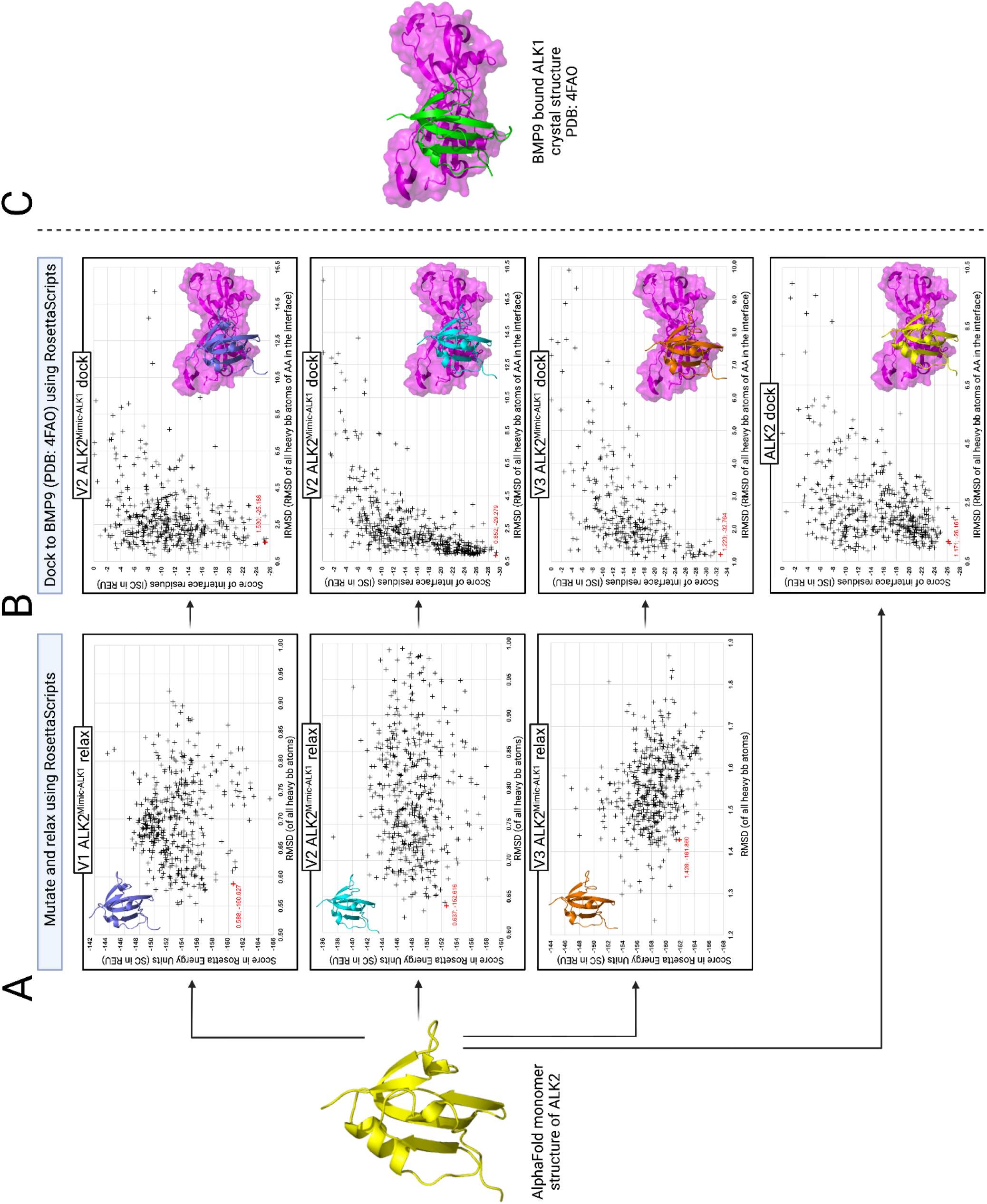
*Supplement to Fig.5* **(A)** Simplified, schematic overview of the *in silico* homology modelling of the variants V1-, V2- and V3 ALK2^Mimic-ALK1^ and docking of all variants and ALK2 to Activin A using the Rosetta Commons Modelling Suite. The basis for the modeling of the variants is the AlphaFold generated structure of ALK2 **(A)** RMSD of all heavy backbone atoms vs. the Rosetta-Score (scoring function ref2015), after introducing mutations and relaxing using the Rosetta Commons Modelling Suite, is plotted for 500 attempts for each variant. The Datapoint of the final homology model (cartoon rep. upper left corner of the plot) is highlighted in red.**(B)** After superimposing all variants and ALK2 to a BMP9 bound ALK1 (PDB: 4FAO) crystal structure and docking using Rosetta, the Interface RMSD (IRMSD) was plotted against the score of the docking interface residues (ISC). Datapoints of the final docked homology models (surface & cartoon representation, lower right corner of the plot) are highlighted in red. **(C)** Cartoon and surface representation of the crystal structure PDB: 4FAO of ALK1 bound to BMP9.

**Supplementary Figure 11:**
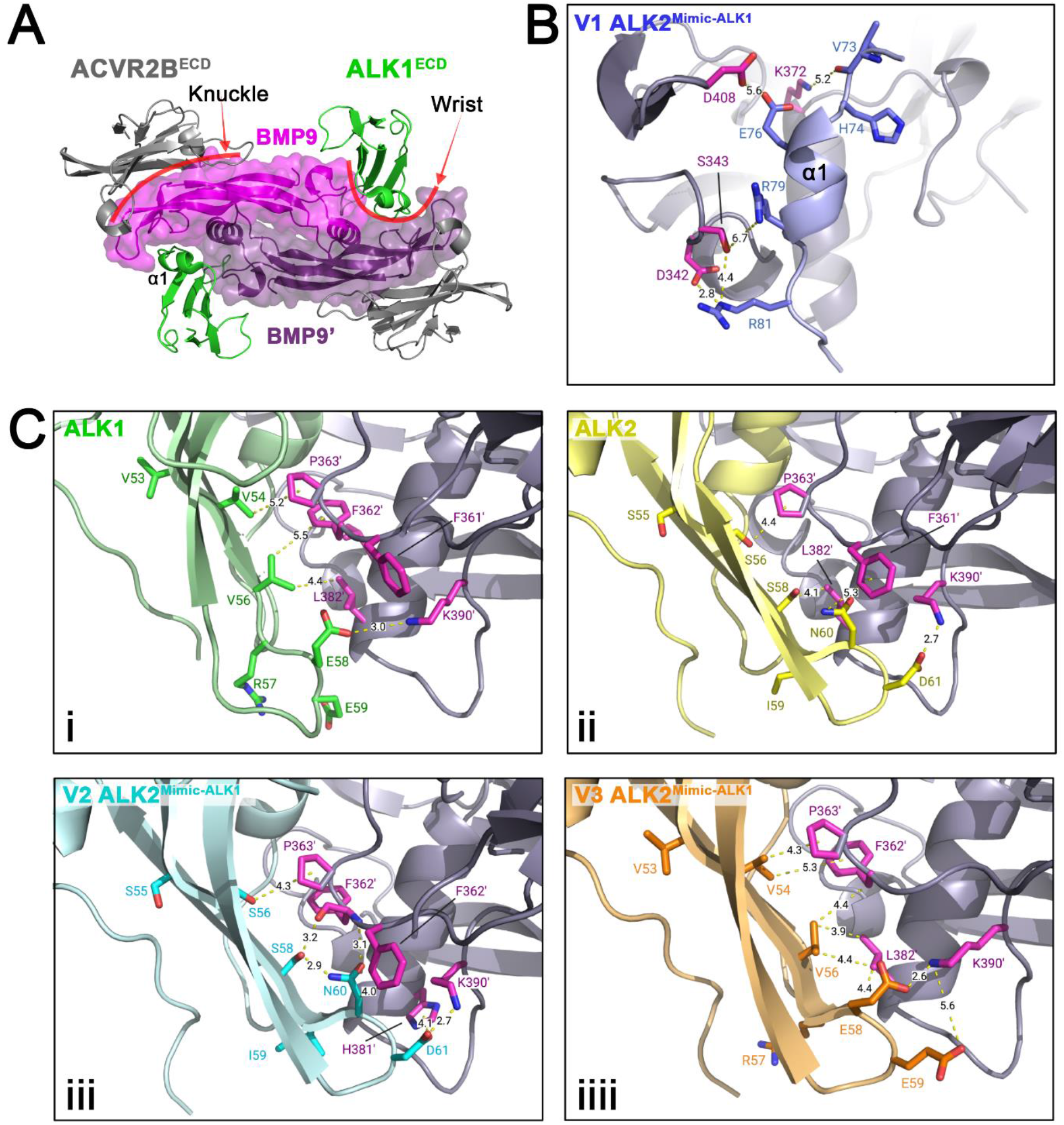
*Supplement to Fig.5* **(A)** Top view of the extracellular domains of ALK1 and ACVR2B in a heterotetrameric receptor complex bound to BMP9 (crystal structure PDB: 4FAO). Type l receptor ALK1 binds at the wrist interface of BMP9. **(B)** HERR”-motif residues of V1 ALK2^Mimic-ALK1^ in cartoon and stick representation. Distance measurements in Angstrom between V1 ALK2^Mimic-ALK1^ and BMP9 were performed in PyMOL **(C)** Residues that were mutated to generate the variant V3 ALK2^Mimic-ALK1^ from V2 ALK2^Mimic-ALK1^ are shown in stick representation in ALK1 (i), ALK2 (ii), V2 ALK2^Mimic-ALK1^ (iii) and V3 ALK2^Mimic-ALK1^ (iiii).

